# Disruption of NOX2-dependent Oxidative Injury with a Targeted Gene-Therapy Approach Prevents Atrial Fibrillation in a Canine Model

**DOI:** 10.1101/765008

**Authors:** Shin Yoo, Anna Pfenniger, Jacob Hoffman, Wenwei Zhang, Jason Ng, Amy Burrell, David A. Johnson, Georg Gussak, Trent Waugh, Suzanne Bull, Brandon Benefield, Bradley P. Knight, Rod Passman, J. Andrew Wasserstrom, Gary L. Aistrup, Rishi Arora

## Abstract

Atrial fibrillation is the most common heart rhythm disorder in adults and a major cause of stroke. Unfortunately, current treatments of AF are suboptimal as they are not targeted to the molecular mechanisms underlying AF. In this study, we demonstrated using a novel gene-based strategy in a clinically relevant large animal of AF that oxidative injury is a key mechanism underlying the onset and perpetuation of AF. First, we demonstrated that generation of oxidative injury in atrial myocytes is a frequency-dependent process, with rapid pacing in canine atrial myocytes inducing oxidative injury through induction of NADPH oxidase 2 (NOX2) and generation of mitochondrial reactive oxygen species. We show that oxidative injury likely contributes to electrical remodeling in AF by upregulating a constitutively active form of acetylcholine-dependent K^+^ current (*I*_KACh_) – called *I*_KH_ - by a mechanism involving frequency-dependent activation of protein kinase C epsilon (PKC_ε_). To understand the mechanism by which oxidative injury promotes the genesis and/or maintenance of AF, we performed targeted injection of NOX2 shRNA in atria of normal dogs followed by rapid atrial pacing. The time to onset of non-sustained AF increased by more than 5-fold in NOX2 shRNA treated dogs. Furthermore, animals treated with NOX2 shRNA did not develop sustained AF for up to 12 weeks. The electrophysiological mechanism underlying AF prevention was prolongation of atrial effective refractory periods, with attenuated activation of PKC_ε_, a likely molecular mechanism underlying this beneficial electrophysiological remodeling. Future optimization of this approach may lead to a novel, mechanism-guided therapy for AF.

**One Sentence Summary:** Targeted disruption of NOX2-dependent oxidative injury with a novel gene therapy approach prevents onset as well as perpetuation of atrial fibrillation.

## Introduction

Atrial fibrillation (AF) is the most common heart rhythm disorder, affecting 2.7 to 6.1 million adults in 2010 in the US alone. AF is a cause of significant morbidity and mortality, being a major cause of stroke (*1*). Because the incidence of AF increases with age, AF is fast becoming an epidemic worldwide due to a rapidly ageing population (*2*). Despite its clinical importance, AF is a difficult condition to treat. Current therapies for AF include anti-arrhythmic drugs, and ablation procedures to electrically isolate the pulmonary veins (*3–5*). Anti-arrhythmic drugs have limited long-term efficacy and can be associated with significant adverse effects, including pro-arrhythmia. Ablation procedures have suboptimal efficacy in the setting of persistent AF, and can be associated with significant complications (*6, 7*). A major reason for the low efficacy of the aforementioned therapies – especially in the setting of persistent AF – is that these therapies are not targeted to the molecular mechanisms underlying the electrical and structural remodeling that is characteristic of long standing, persistent AF (*8*). A better understanding of the molecular mechanisms underlying AF is essential for the development of innovative and improved therapeutic approaches for this condition.

Oxidative injury, which is an imbalance between generation and neutralization of reactive oxygen species (ROS), is thought be a major mechanism underlying AF and is regarded as a possible therapeutic target for this condition (*9–12*). Inflammation and oxidative injury have been described in atrial appendages from patients with AF (*13*). Furthermore, redox potentials of glutathione have been associated with increased prevalence and incidence of AF(*14*). ROS generated in the cardiovascular system are primarily derived from NADPH oxidases (NOX), mitochondrial electrical transport chain, xanthine oxidase and uncoupled eNOS (*15, 16*). NADPH oxidase 2 (NOX2) has been suggested to be a major source of oxidative injury in atrial appendages of patients with AF. Moreover, mice with cardiac-specific overexpression of Rac-1, a necessary activator for NOX2, develop AF spontaneously (*17*), supporting a role for NOX2 in determining an atrial substrate for the new onset of AF. More recent studies suggest that with increasing duration of AF, there is an increase in not only expression of NOX2 but also mitochondrial ROS in atrial tissue (*18*). Although these studies suggest a likely role for oxidative injury in the genesis and/or maintenance of AF, the precise mechanisms by which oxidative injury creates a vulnerable substrate for AF in the *intact* atrium are not known. Specifically, it is not known which atrial ion channels involved in electrical remodeling in AF are most vulnerable to oxidative injury, and whether this oxidative injury actually leads to effective refractory period (ERP) shortening in the intact atrium. Indeed, although a number of ion channels and transporters including L-type Ca^2+^ channel, Na^+^ channel, transient outward K^+^ channel and type 2 ryanodine receptor (RyR2) have been shown to be redox-sensitive (*19*), only a few studies have looked at this in the context of AF (*20*).

Among the ion channels that have been suggested to play a key role in action potential duration (APD) and ERP shortening in AF, the best studied are the L-type Ca^2+^ current (*I*_CaL_) – which is downregulated in AF – and the inward-rectifier K^+^ current (*I*_K1_), which is elevated in AF. Lately, several studies have described a form of the acetylcholine-dependent K^+^ current (*I*_KACh_) that is constitutively active. This current -called *I*_KH_ - has been invoked in ERP shortening in the rapid atrial pacing (RAP) model, as well as in patients with paroxysmal and persistent AF (*21, 22*). However, the precise mechanisms underlying the emergence of *I*_KH_ in the setting of AF are not known. What is known is that this current is PKC sensitive, with the conventional PKC isoform PKC_α_ likely involved in inhibition of this current, but with the novel PKC isoform PKC_ɛ_ acting to stimulate the emergence of *I*_KH_ (*23*). This PKC_ɛ_ mediated increase in *I*_KH_ was demonstrated to be a frequency-dependent phenomenon, with increasing frequency of atrial tachypacing leading to membrane translation of PKC_ɛ_ and an increase in magnitude of *I*_KH_. Because PKC isoforms are well known to be acute phase reactants in the heart, with PKC_ɛ_ having been shown to be highly sensitive to stressors such as oxidative injury (*24, 25*), we hypothesized that upregulation of *I*_KH_ in RAP-induced AF is mediated by a frequency-dependent increase in oxidative injury, with resulting activation of PKC_ɛ_. In order to determine the precise role of oxidative injury in causing electrical remodeling in the intact atrium, we further hypothesized that oxidative injury leads to not only the initiation but also the maintenance of ERP shortening in the intact, fibrillating atrium. To examine this hypothesis, we performed targeted expression of anti-NOX2 short hairpin RNA (NOX2 shRNA) in the intact atria of dogs, and then subjected these animals to RAP for a period of several weeks. Using this novel gene therapy approach, we demonstrated a pivotal role for NOX2-generated oxidative injury in the creation as well as the maintenance of electrical remodeling in AF. The results of this study yield valuable mechanistic insights into the pathogenesis of AF and have important therapeutic implications for the clinical management of this common arrhythmia.

## Results

### ROS generation in atrial myocytes is a frequency-dependent phenomenon

Electrical remodelling in the AF atrium is largely thought to be the result of rapid atrial rates, with rapid stimulation leading to ion channel remodelling such as a decrease in *I*_CaL_. More recently, frequency-dependent *I*_KH_ upregulation has been demonstrated in atrial myocytes. The upstream signalling mechanisms underlying this ion channel remodelling are not well understood. Since we believe that oxidative injury is a major mechanism underlying electrical remodelling in AF, we hypothesized that ROS generation in atrial myocytes is a frequency-dependent phenomenon, with increasing stimulation frequency leading to progressively increasing ROS. We therefore investigated ROS generation in isolated-paced atrial swine myocytes in response to increasing pacing frequency. Total cellular fluorescence from a ROS sensitive indicator, CellROX Deep Red, increased in a frequency dependent manner, increasing progressively from control (unpaced) to 2 Hz (Figure 1A). These results support our postulate that ROS generation in atrial myocytes is a frequency-dependent phenomenon.

**Fig. 1.**
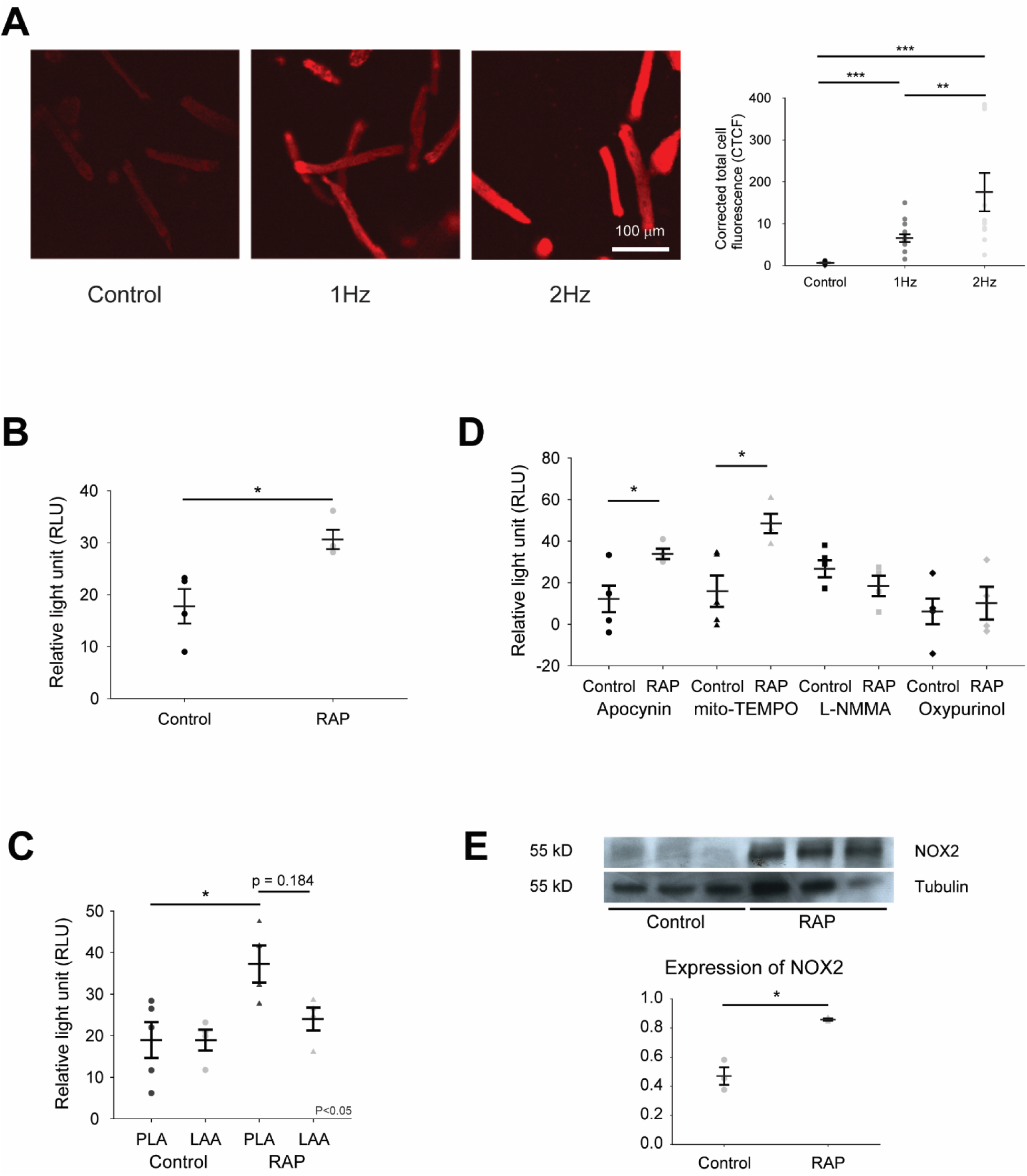
Frequency-dependent ROS generation in isolated pig myocytes and in canine RAP left atrium. (A) ROS generation in tachypaced swine atrial myocytes. (B) Superoxide generation in control and RAP left atrium. (C) Superoxide generation in PLA versus LAA, in RAP and control atria. (D) Relative contribution of different enzymatic sources of ROS to superoxide generation in RAP left atrium. (E) Immunoblot and densitometric measurements of NOX2 (normalized to tubulin) from control and RAP atria. Data are presented as mean ± SEM; * p < 0.05, ** p < 0.01 and *** p < 0.001.

### NOX2 and mitochondria-generated superoxide (O_2_^-^) significantly increases in rapidly paced left atrium

Next, we looked for evidence of oxidative injury in the intact, fibrillating atrium. Lucigenin chemiluminescence assay revealed a significant increase in overall O_2_^-^ generation in left atrial tissue homogenate from RAP dogs as compared to control (Figure 1B). While O_2_^-^ generation increased in both the posterior left atrium (PLA) and left atrial appendage (LAA) in RAP atria, the increase was significant only in the PLA (Figure 1C). Next, we determined relative contribution to O_2_^-^ generation by various enzymatic sources of ROS by the application of specific ROS inhibitors. There was higher activity of mitochondrial ROS and NADPH oxidase (NOX2) in RAP PLA, compared to control PLA (Figure 1D). Consistent with the increase in activity of NOX2 in RAP PLA, expression of the gp91 subunit of NOX2 was also significantly greater in RAP, as compared to control atria (Figure 1E).

### I_KH_ is highly sensitive to ROS inhibition

Since ROS are elevated in the rapidly paced atrium, and since *I*_KH_ - which is thought to contribute to ERP shortening in the fibrillating atrium – is stimulated by the acute phase reactant PKC_ɛ_, we hypothesized that *I*_KH_ induction in RAP is modulated by ROS. We therefore investigated the effect of various inhibitors of mitochondrial ROS and NOX2 (the two sources of ROS found to be elevated in the rapidly paced atrium) on *I*_KH_, *I*_KACh_, and the inward rectifier K current *I*_K1_. *I*_KH_ and *I*_KACh_ were measured as the Tertiapin Q- and Carbachol (CCh) - sensitive current, respectively by calculating differences between the currents before and after the application of drug. *I*_K1_ was referred as residual inward current after TQ.

### I_KH_ is attenuated by mitochondrial ROS inhibition

We first examined the effect of mitochondrial ROS inhibition in RAP myocytes using mito-TEMPO on *I*_KACh_. Figure 2A shows currents elicited by 4 second step pulses from a holding potential of -40 mV to voltage between -120 mV and -20 mV in control and 50 μM mito-TEMPO treated rapidly paced left atrial myocytes. *I*_KACh_ refers to carbachol (CCh) sensitive current calculated by subtraction of currents before and after the application of 10 μM CCh. Representative current traces (left and middle panels) and mean current-voltage relations (I-V curve, right panel) demonstrate that there is no significant difference in *I*_KACh_ between control and mito-TEMPO pre-incubated RAP myocytes.

**Fig. 2.**
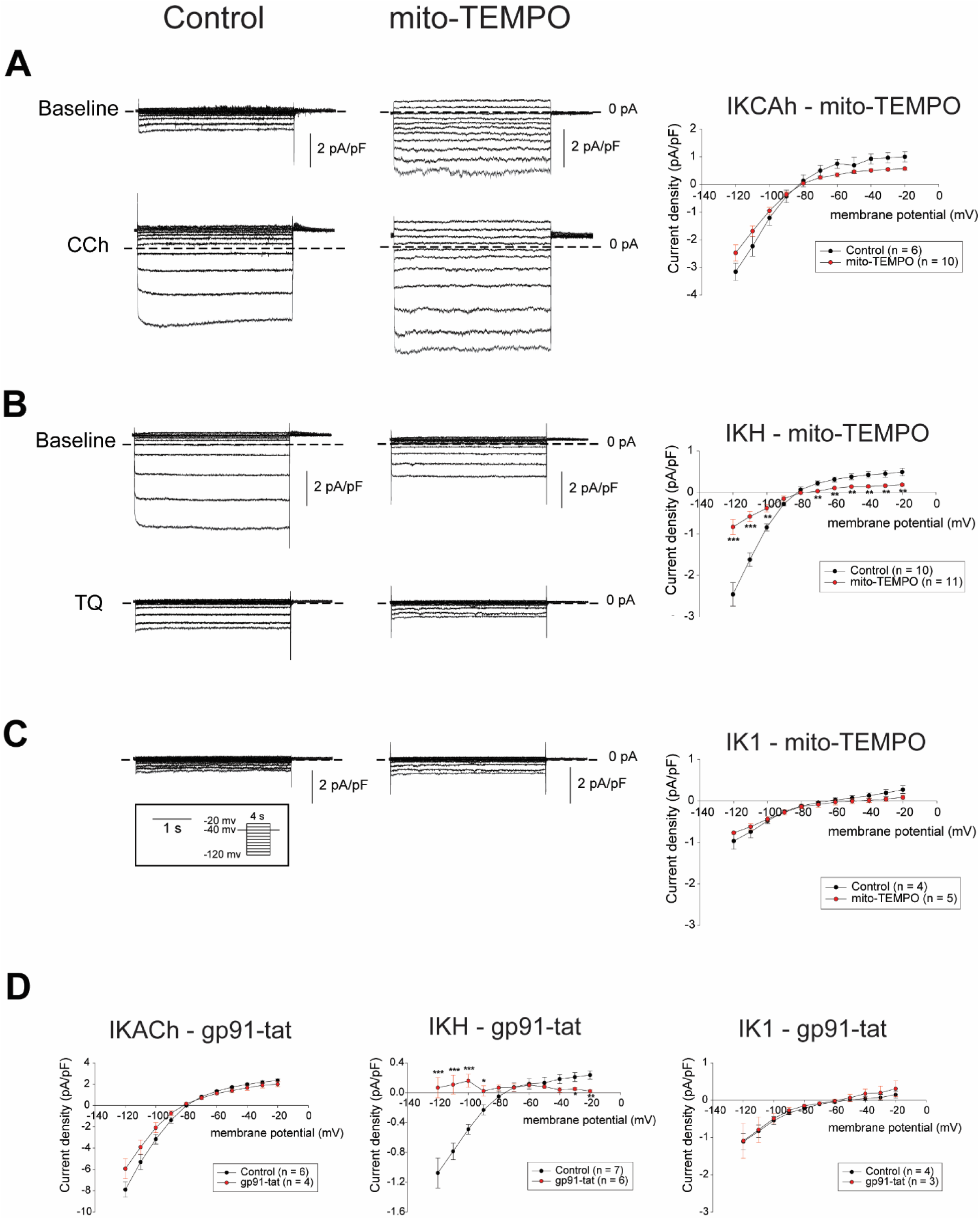
Effect of mito-TEMPO and gp91-tat on inwardly rectifying currents in RAP atrial myocytes. (A - C) Raw traces (left and middle panels) of *I*_KACh_ (A), *I*_KH_ (B) and *I*_K1_ (C) elicited by 4 seconds step pulses from a holding potential of -40 mV to voltage between -120 mV and – 20 mV(pulse protocol shown in inset) and I-V curve (right panels) for control and mito-TEMPO pre-incubated RAP atrial myocytes. (D) I-V curve of *I*_KACh_ (left), *I*_KH_ (middle) and *I*_K1_ (right) for control and in the presence of gp91-tat elicited by 400 ms ramp pulses from a holding potential of -40 mV to voltage between 20 mV and -120 mV. Data in I-V plots are presented as mean ± SEM at given membrane potentials; * p < 0.05, ** p < 0.01 and *** p < 0.001.

We then assessed the effect of mitochondrial ROS inhibition on *I*_KH_. Figure 2B shows representative recordings and mean I-V curve for the currents elicited by the same step pulse protocol as *I*_KACh_. *I*_KH_ refers to Tertiapin Q (TQ) – a selective blocker for *I*_KH_ – sensitive current calculated by subtraction of currents before and after application of 100 nM TQ. Representative current traces (left and middle panels) and mean I-V curve (right panel) demonstrate that *I*_KH_ was significantly attenuated in mito-TEMPO pre-incubated atrial myocytes, compared to control RAP myocytes.

Next, we determined the effect of mitochondrial ROS inhibition on the inward rectifier current *I*K1, calculated as the residual current elicited by same step pulse protocol after TQ application. Figure 2C shows representative traces of *I*_K1_ (left and middle panels) and mean I-V curve (right panel). These results demonstrate that there is no significant difference in *I*_K1_ in mito-TEMPO pre-incubated RAP myocytes, compared to control RAP myocytes.

Lastly, since *I*_CaL_ is known to be downregulated in AF atria and is thought to play an important role in ERP shortening in AF (*2, 26*), we also examined the effect of mitochondrial ROS inhibition on *I*_CaL_. Consistent with the literature, our own data demonstrated a significant reduction of *I*_CaL_ in RAP myocytes, compared to normal atrial myocytes (Supplementary Figure S1). As shown in Supplementary Figure S1, pre-incubation with mito-TEMPO did not lead to any significant change in in *I*_CaL_ in RAP myocytes.

Taken together, these results indicate that of the major ion channels thought to contribute to ERP shortening in the rapidly paced atrium, *I*KH was the only current sensitive to mitochondrial ROS inhibition.

### I_KH_ is attenuated by NOX2 inhibition

Next, we examined the effect of NOX2 inhibition on *I*_KACh_, *I*_KH_ and *I*_K1_ in isolated RAP myocytes. We examined currents elicited by 400 millisecond (ms) ramp pulses from a holding potential of -40 mV to voltage between 10 mV and -120 mV in control condition and in the presence of 50 μM of gp91-tat, which is a specific NOX2 inhibitory peptide, in the pipette solution in right atrial myocytes. The mean I-V curve (left panel) demonstrate that there is no significant difference in *I*_KACh_ between control and gp91-tat treated RAP myocytes (Figure 2D). Of note, the average amplitude of *I*_KACh_ in right atrial myocytes was noted to be 2 fold larger than in left atrial myocytes (Figure 2A), consistent with prior studies (*27*).

We next examined the effect of gp91-tat on *I*_KH_. Figure 2D (middle panel) shows the mean I-V curve for *I*_KH_ elicited by same ramp pulse protocol as *I*_KACh_ in right atrial myocytes. These results demonstrate that *I*_KH_ was significantly attenuated in the presence of gp91-tat in the pipette solution, compared to control conditions.

We also examined the effect of gp91-tat on *I*_K1_. Figure 2D (right panel) shows the mean I-V curve in right atrial myocytes. These results demonstrate there is no significant difference in *I*_K1_ in the presence of gp91-tat, compared to control condition.

Taken together, *I*_KH_ was the only inwardly rectifying K current significantly affected by NOX2 inhibition. In view of NOX2 being a major enzymatic source of ROS in the rapidly paced atria (Figure 1D), the clear attenuation of *I*_KH_ at voltages from -90 mV through -110 mV in the presence of gp91-tat supports a likely role for NOX2 in the emergence of *I*_KH_ in RAP myocytes.

### ROS-induced increase in I_KH_ during RAP is mediated by enhanced PKCε signalling

Previous studies have demonstrated that enhanced PKC signalling may be contributing to the emergence of *I*_KH_ in RAP myocytes as well as in myocytes from patients with AF (*21, 23*). Furthermore, Makary et al. showed that membrane translocation of the stimulatory PKC isoform PKC_ɛ_ may be essential to the emergence of *I*_KH_ in canine RAP myocytes (*23*). Since PKC signalling and the isoform PKCε have been shown to be acute phase reactants (*24, 25*), we hypothesized that ROS-mediated emergence of *I*_KH_ in RAP myocytes is at least partially mediated by an increase in PKC – and specifically, PKC_ɛ_ - signalling.

We therefore first examined the effect of bisindolylmaleimide-1 (BIM1) – a non-specific PKC inhibitor – on *I*_KH_ in the absence and presence of mito-TEMPO by whole cell patch clamp via eliciting 4 s step pulses. As anticipated, *I*_KH_ was attenuated by the application of 2 μM of BIM1, as compared to control (Supplementary Figure S2A). In mito-TEMPO pre-incubated myocytes, which exhibited around 70% attenuation of *I*_KH_ (Figure 2B), BIM1 had no additional effect on *I*_KH_ (Supplementary figure S2B). These results indicate that BIM1 effect on *I*_KH_ may be at least partially mediated by ROS. Similarly, a specific PKC_ɛ_ inhibitory peptide was also found to have no synergistic effect on mito-TEMPO induced attenuation of *I*_KH_ in RAP myocytes (Supplementary figure S2C).

Since PKC_ɛ_ is an acute phase reactant (*24, 25*), we further hypothesized ROS is upstream of PKC_ɛ_ signalling in the rapidly paced atrium. We therefore assessed membrane translocation of PKC_ɛ_ in isolated atrial myocytes from normal dogs subjected to *in-vitro* tachypacing, in the absence and presence of the non-specific ROS inhibitor, N-acetylcysteine (NAC). Consistent with the previous report by Makary et al. (*23*), we discovered that pacing of isolated atrial myocytes at 3 Hz compared to 1 Hz led to increased membrane translocation of PKC_ɛ_ (Figure 3A and 3B). However, in the presence of 10 mM NAC, the membrane translation of PKC_ɛ_ induced by 3Hz tachypacing was significantly attenuated (Figure 3A and 3C). These aforementioned data indicate that ROS induced increase in *I*_KH_ in the rapidly paced atrium is at least partially mediated by increased PKC_ɛ_ signalling.

**Fig. 3.**
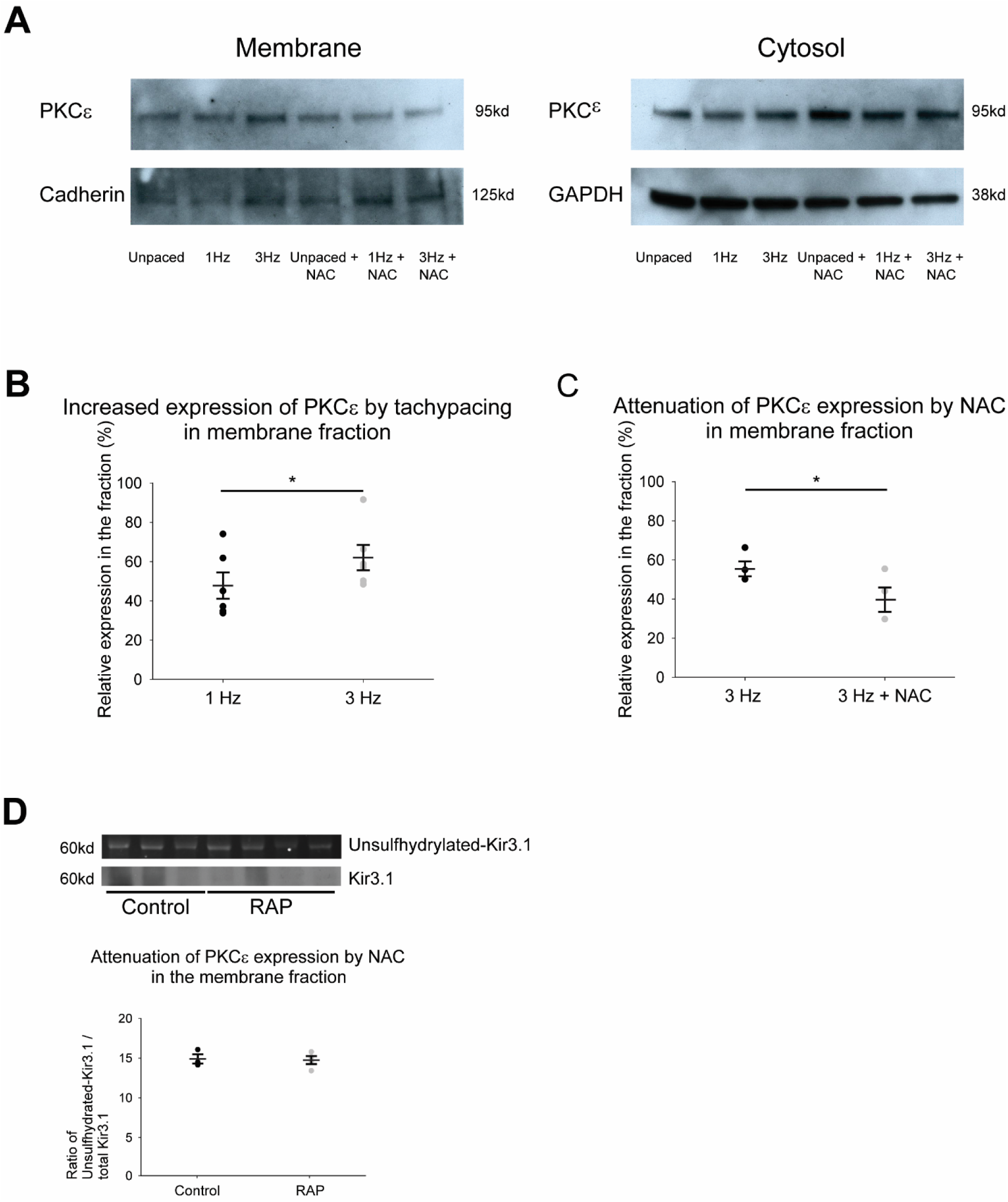
ROS inhibition attenuates tachypacing-induced membrane translocation of PKC_ɛ_ in atrial myocytes. (A) Representative immunoblot for PKC_ɛ_ in membrane and cytosolic fractions, along with GAPDH and Cadherin as loading controls. (B and C) Densitometric measurements of PKC_ɛ_ in membrane fraction after *in vitro* tachypacing at 1 Hz, 3 Hz and 3 Hz with NAC (D) Top band, unsulfhydrylated-Kir3.1; bottom band, immunoblot for Kir3.1. Densitometric measurements showed level of unsulfhydrylated-Kir3.1 (normalized to Kir3.1). Data are presented as mean ± SEM; * p < 0.05.

To determine whether direct oxidation of the Kir3.1/Kir3.4 which encode *I*_KACh_ channel may also be playing a role in the emergence of *I*_KH_, we assessed glutathionylation level of Kir3.1 in rapidly paced versus control atrium. As shown in figure 3D, there was no significant increase in oxidation (glutathionylation) of Kir3.1 in RAP as compared to control atrium.

### AF inducibility and duration is markedly reduced after NOX2 shRNA treatment

Since NOX2 appears to be a major contributor to ROS generation in rapidly paced atria, and since NOX2 inhibition was noted to attenuate *I*_KH_ in RAP myocytes, we hypothesized that targeted inhibition of NOX2 in the atria would prevent RAP-induced ERP shortening and consequent AF. We elected to selectively target NOX2 in the atrium by using NOX2 shRNA. We generated a shRNA to canine NOX2 (see Supplementary figure S3A for target sequence). Using this shRNA, we were able to achieve significant knockdown of NOX2 in HEK 293 cells (Supplementary figure S3B).

Six dogs underwent sub-epicardial injection of NOX2 shRNA in the atria, followed by electroporation to facilitate myocardial gene transfer. Gene injection and electroporation was limited to the PLA in the first 4 animals, with subsequent two animals receiving gene injection in the left atrial free wall (LAFW), LAA, and right atrium as well. Eighteen animals receiving either injection of scrambled shRNA or no gene injection were used as control. See Figure 4 for experimental design for assessment of AF both in the short term (i.e. 4 weeks of RAP) and in the long term (i.e. 12 weeks of RAP). After gene injection, animals were subjected to RAP and duration of induced AF was subsequently recorded during periods in which RAP was interrupted. Figure 5 shows the duration of AF after initiation of RAP. As detailed in panel A, whereas control animals developed sustained AF for more than 30 minutes within a median of 4 days of RAP (interquartile range (IQR) 4-9 days), it took 21 days or longer for NOX2 shRNA animals to develop this burden of AF (p<0.01). Three animals in each group were followed for twelve weeks to assess development of persistent AF (defined as AF duration longer than 8 hours; also see Methods). Panel B shows that it took a median of 14 days for control animals to develop >8 hours of AF. In striking contrast, no animal receiving NOX2 shRNA developed AF>8 hours, despite up to 12 weeks of RAP (p<0.05). Over the entire recorded period, control animals spent a median of 60 minutes in AF (IQR 30-60 minutes), whereas NOX2 shRNA animals spent a median of 0 minutes in AF (IQR 0-2 minutes) (p = 0.008). Representative examples of intra-cardiac electrograms are shown in Figure 5C. The dominant rhythm in NOX2 shRNA animals was either sinus rhythm (top right) or atrial flutter (top left and top middle). In distinct contrast, AF was the dominant rhythm in all control animals (bottom panels).

**Fig. 4.**
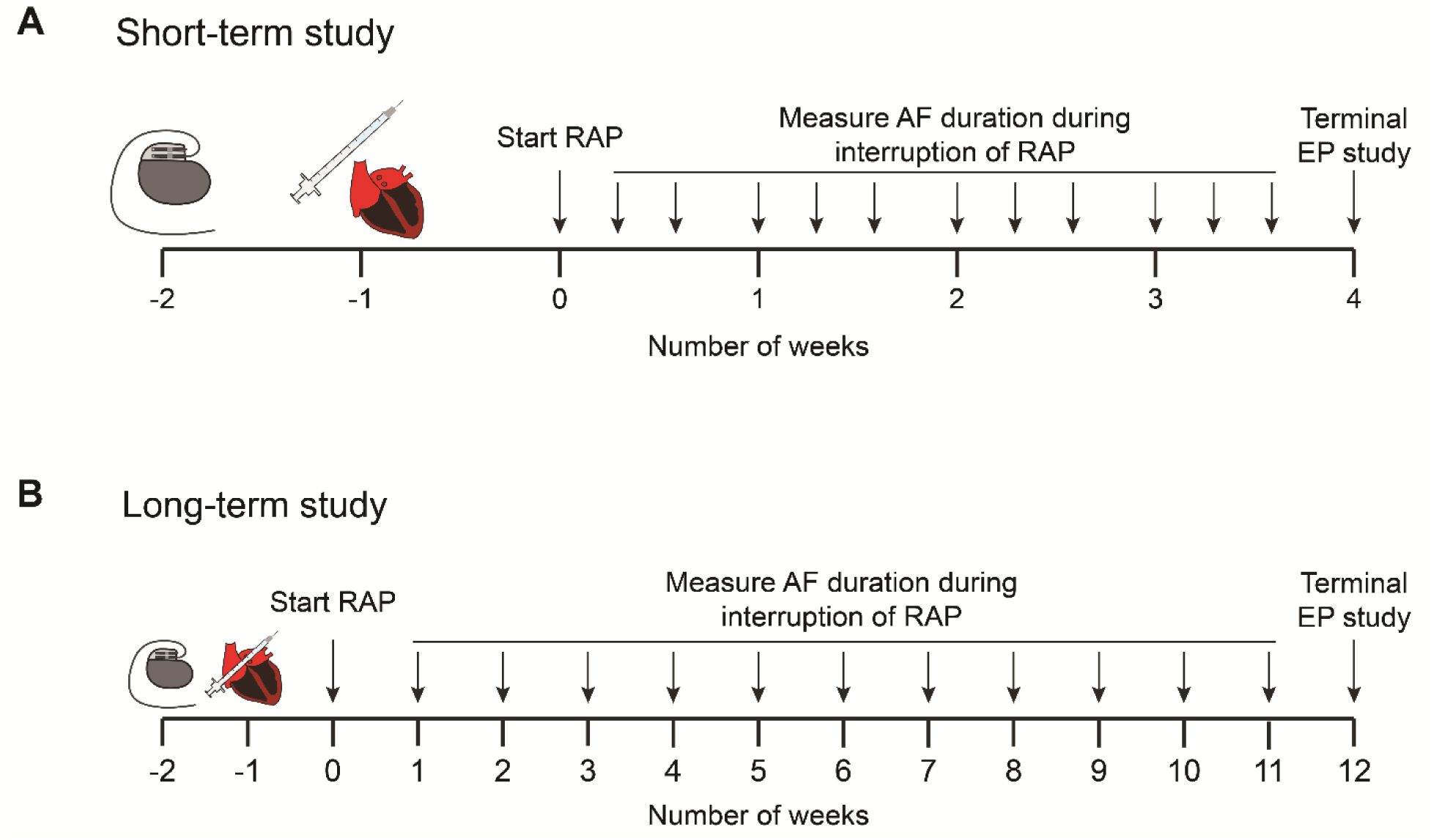
Experimental design. One week after pacemaker implantation, animals underwent open-chest subepicardial gene injection in the atria, followed by electroporation. After one week of rest, RAP was initiated. RAP was interrupted 3 times a week for 30 minutes for the first 28 days (top panel). For the long-term study, some animals were followed for 12 weeks, and RAP interrupted weekly for up to 8 hours (bottom panel).

**Fig. 5.**
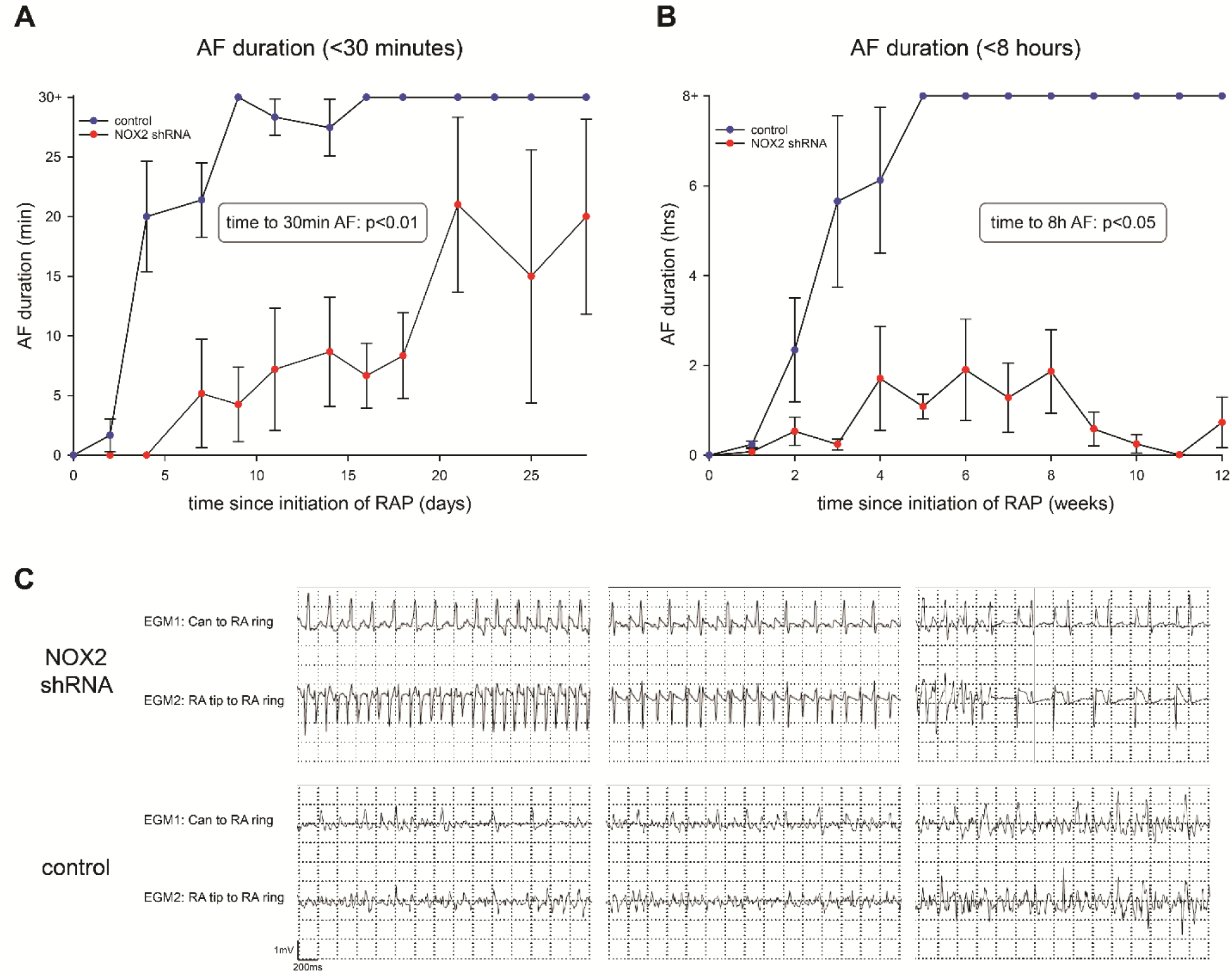
NOX2 shRNA prevents development of sustained AF. (A) Animals who received NOX2 shRNA developed significantly shorter AF, with a delay in development of sustained AF > 30 minutes. N = 3-12 for controls, n = 3-5 for NOX2 shRNA. (B) NOX2 shRNA gene injection prevented development of sustained AF > 8 hours. (C) Representative examples of intracardiac EGMs are shown for NOX2 shRNA animals (top) and control (bottom). Corresponding Can to RA ring and RA tip to RA ring EGMs are shown, with evidence of atrial flutter or sinus rhythm for NOX2 shRNA, and AF for control. Data are mean ± SEM; * p<0.05, ** p<0.01, *** p<0.001.

### Left atrial ERPs are prolonged after NOX2 shRNA treatment

At the terminal study, atrial ERPs were measured in the PLA and LAA in animals that were in sinus rhythm or in which sinus rhythm was restored by burst pacing or electrical cardioversion. As shown in Figure 6A, combined ERPs were significantly longer in NOX2 shRNA injected dogs as compared to controls. This ERP lengthening was noted in the PLA as well as the LAA.

**Fig. 6.**
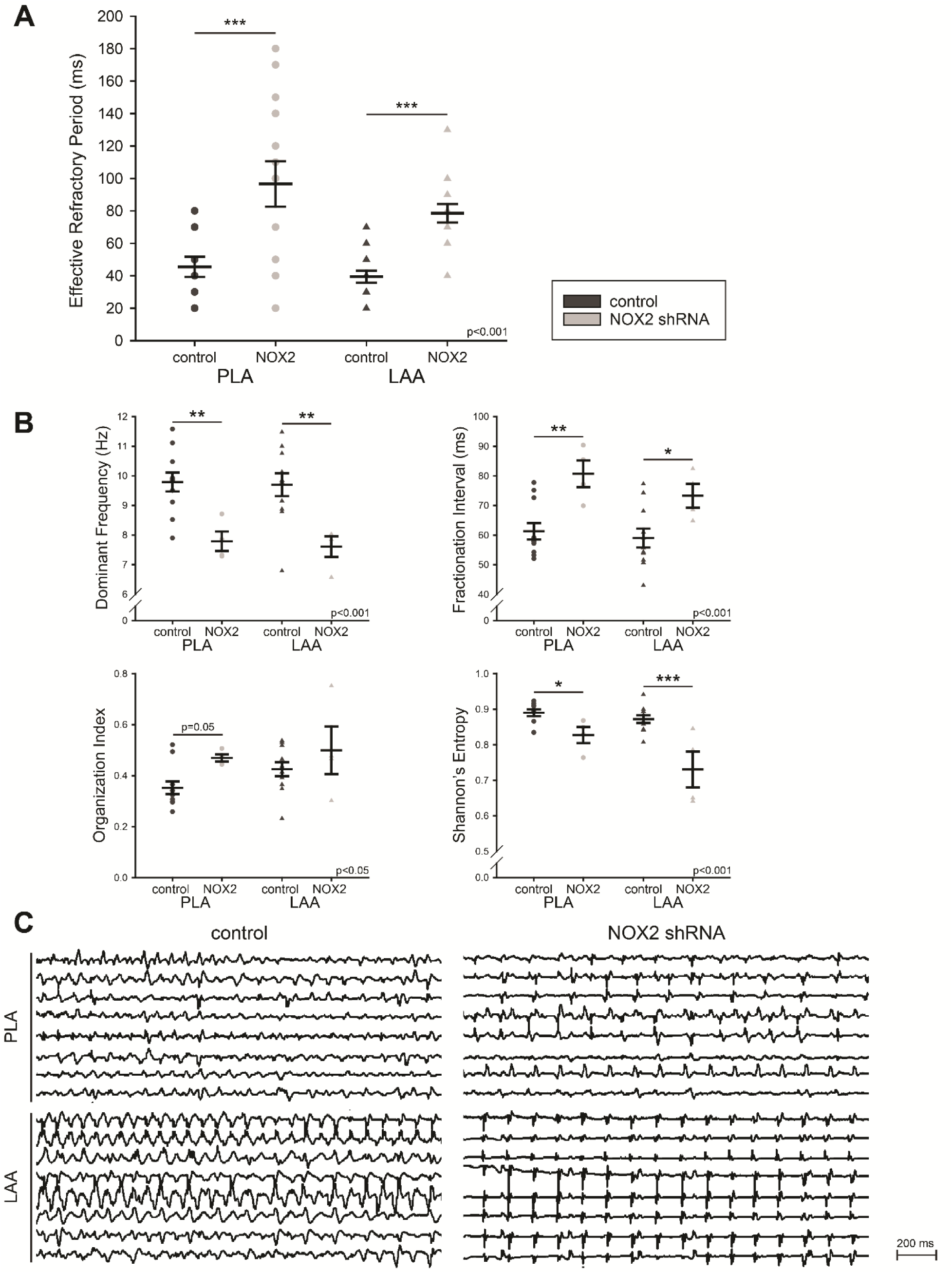
NOX2 shRNA prevents electrical remodeling induced by RAP. (A) ERPs were measured in the posterior left atrium (PLA) and left atrial appendage (LAA) of 6 control and 4 NOX2 shRNA animals after RAP. Results are shown as mean ± SEM of combined ERPs. (B). AF electrograms are slower and more organized after NOX2 shRNA gene injection (n=4) compared to control (n=11) with higher dominant frequency, longer fractionation interval, higher organization index and lower Shannon’s Entropy. Data are mean ± SEM; * p<0.05, ** p<0.01, ***p<0.001. (C) Representative electrograms of control (left) and NOX2 shRNA animals (right) in the PLA (top) and LAA (bottom).

### Residual AF after NOX2 shRNA treatment is slower, more organized and less complex than AF in controls

Periods of AF were recorded during the terminal study, either when the animals were spontaneously in AF, or after AF induction with burst pacing. AF in NOX2 shRNA animals showed a decrease in DF (a frequency domain measure of activation rate), an increase in FI (the mean interval between deflections detected in the electrogram segment), an increase in OI (a frequency domain measure of temporal organization or regularity), and a decrease in ShEn (a statistical measure of complexity) when compared to controls (Figure 6B and 6C). Taken together, these data demonstrate the limited AF that could be induced in NOX2 shRNA animals was significantly slower and more organized than the AF noted in control animals.

### NOX2 shRNA treatment does not prevent RAP induced changes in ventricular function

A transthoracic echocardiogram was performed at baseline and prior to the terminal study in 5 control animals and 3 NOX2 shRNA animals. Supplementary Table S1 details our findings. There was no significant difference in any echocardiographic parameters between groups at baseline. RAP caused a reduction in left ventricular ejection fraction (LVEF) and left ventricular (LV) global longitudinal strain in both groups, without significant difference between animals having received NOX2 shRNA and controls. Right ventricular (RV) systolic function also appeared significantly reduced in control animals as determined by tricuspid annular plane systolic excursion by M mode (TAPSE) and RV s’ velocity, with a similar trend in NOX2 shRNA animals. Importantly, there was no difference in left atrium size and left atrium reservoir strain between baseline and after RAP, and between groups.

### NOX2 is attenuated in NOX2 shRNA injected atria

After the terminal electrophysiological study, atrial tissue was removed to assess for gene expression, oxidative injury and effect of gene on signaling. As shown in Figure 7A, dogs that underwent NOX2 shRNA injection demonstrated >50% decrease in native NOX2 expression in gene-injected PLA when compared to the same region in control animals, and also when compared to a non-injected region (i.e. LAA) as shown in Figure 7B. This decrease in NOX2 mRNA was accompanied by a decrease in NOX2 protein expression in the PLA (Figure 7C).

**Fig. 7.**
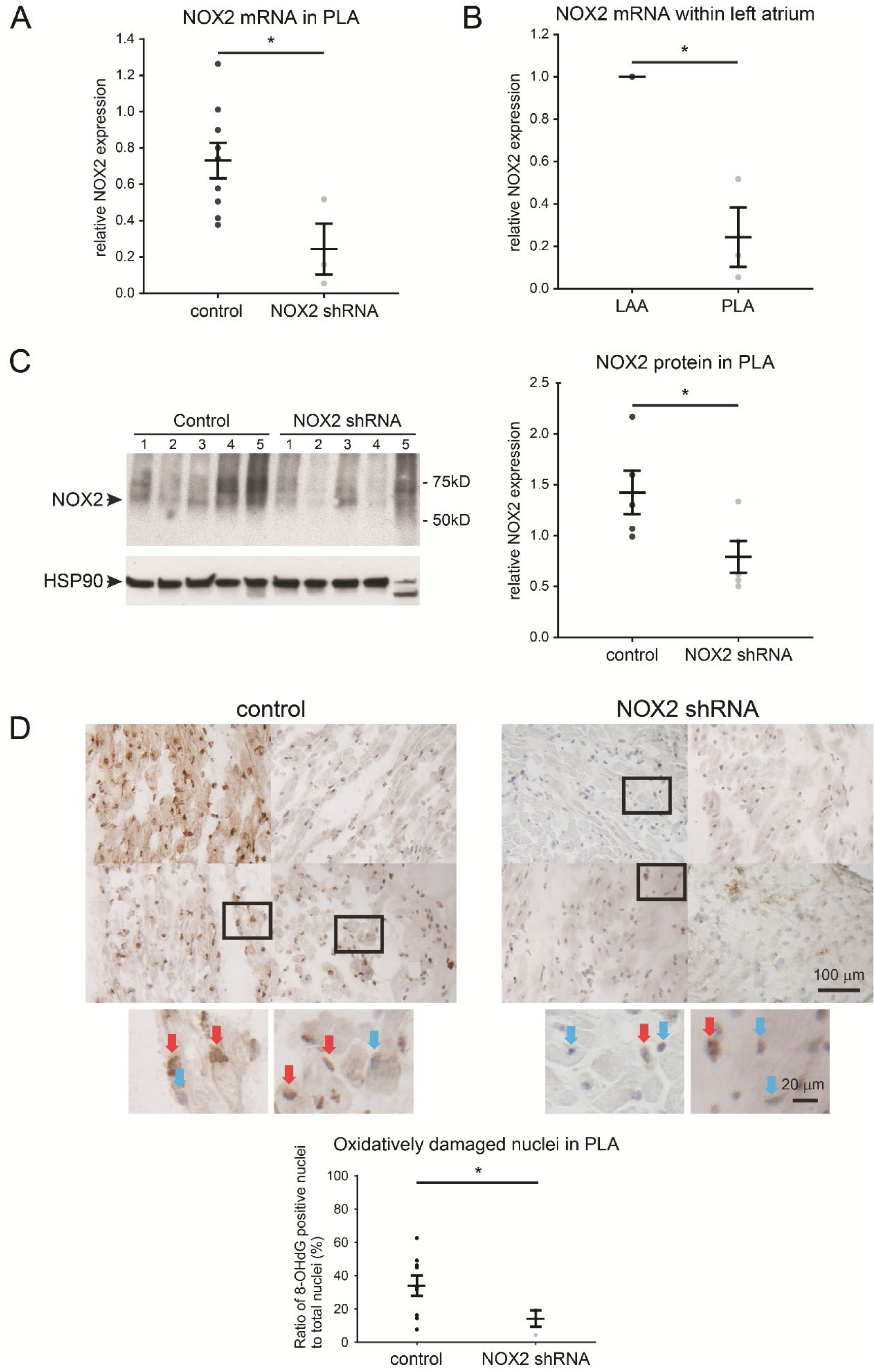
NOX2 shRNA attenuates NOX2 levels and attenuates DNA oxidative damage in RAP atrium. (A) NOX2 qPCR in animals injected in the posterior left atrium (PLA) alone (n = 3) shows effective gene suppression when compared to controls PLA (n=9) and (B) when compared to an uninjected left atrial region (left atrial appendage, LAA, n=3). All samples were normalized to respective LAA. (C) Protein expression of NOX2 is attenuated by injection of NOX2 shRNA (n=5 in each group). (D) 8-OHdG stained tissue sections in control RAP PLA (n = 9) and NOX2 shRNA transfected PLA (n = 3). 8-OHdG positive nuclei were stained in brown and indicate oxidatively damaged nuclei. Blue stained nuclei indicate undamaged nuclei. Inset; undamaged and oxidatively damaged nuclei are designated by blue and red arrows, respectively. Ratio of 8-OHdG positive nuclei against total number of nuclei is shown on the left. N=9 for control and n=3 for NOX2 shRNA. Data are presented as mean ± SEM; * p< 0.05.

### DNA oxidative damage is attenuated by NOX2 shRNA

To determine whether the decrease in native NOX2 by NOX2 shRNA was accompanied by a decrease in oxidative damage in the atrium, we examined the levels of 8-hydroxy-2-deoxyguanosine (8-OHdG) - a biomarker of oxidative damage of DNA – in gene-transfected PLA. Figure 7D demonstrates significant attenuation in the percentage of oxidatively damaged nuclei (i.e. 8-OHdG stained nuclei) in NOX2 shRNA compared to control dogs.

### RAP-induced membrane translocation of PKC_ε_ is attenuated by NOX2 shRNA treatment

As had been performed in isolated atrial myocytes, we also determined PKC_ε_ expression in separated cytosolic and membrane protein fractions of left atrial tissue from animals subjected to RAP. Similar to a previous report (*23*), we discovered that RAP increased the relative membrane expression of PKC_ε_, as compared to controls, consistent with a RAP-dependent translocation of PKC_ε_ from the cytosol (Figure 8A).

**Fig. 8.**
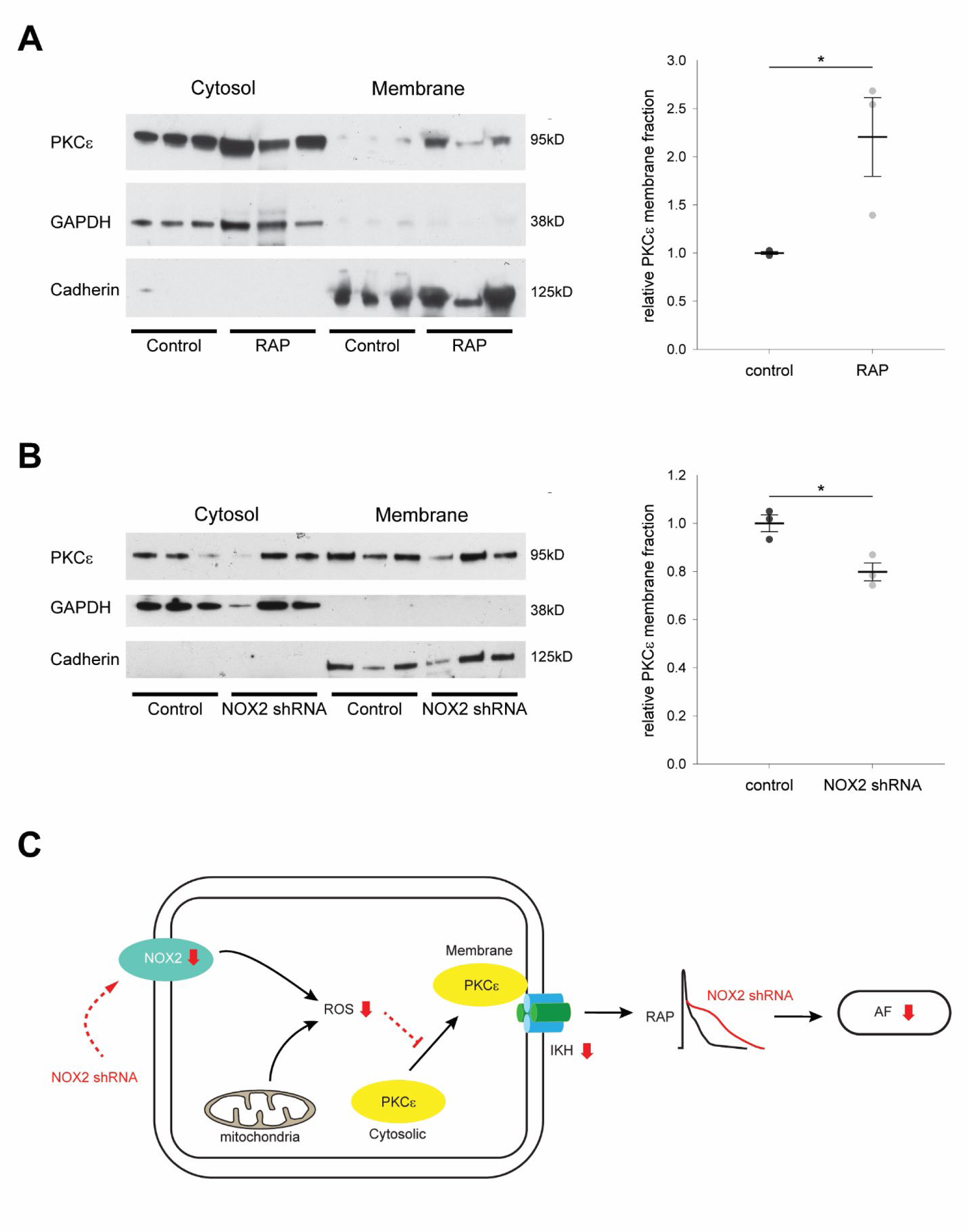
Rapid atrial pacing (RAP) causes membrane translocation of PKC_ε_, which is inhibited after NOX2 shRNA gene injection. (A) Western blots of atrial tissue from 3 control (C) and 3 RAP animals for PKCε, GAPDH as cytosol loading control, and Cadherin as membrane loading control are shown (left panel). Mean ± SEM of relative membrane fraction of PKCε in control and RAP animals (right panel). (B) Western blots of atrial tissue from 3 control (C) and 3 NOX2 shRNA animals (NOX2) after RAP for PKCε, GAPDH as cytosol loading control, and Cadherin as membrane loading control are shown (left panel). Mean ± SEM of relative membrane fraction of PKCε in control and NOX2 shRNA animals (right panel). * p<0.05. (C) Schematic illustration of potential mechanisms by which NOX2 shRNA transfection prevents electrical remodeling in AF.

We therefore also assessed the effect of NOX2 shRNA on membrane translocation of PKC_ε_ in intact atrium. When compared to control animals, relative membrane expression of PKC_ε_ was reduced in dogs receiving NOX2 shRNA (Figure 8B). This demonstrates that NOX2 shRNA attenuates RAP-dependent translocation of PKC_ε_ from the cytosol to the membrane.

### Expression of ion channels involved in ERP shortening during RAP

We evaluated left atrial gene mRNA expression of ion channels thought to underlie ERP shortening in AF, in animals that received NOX2 shRNA and controls. Supplementary Figure 4 summarizes our qPCR results. Expression of KCNJ2, KCNJ3, and KCNJ5 did not differ significantly in the left atrium of NOX2 shRNA animals when compared to controls. Overall ANOVA was significant when evaluating KCNJ12 expression, but pairwise comparisons between conditions or left atrial regions did not reach significance. CACNA1C was significantly higher in the control PLA when compared to control LAA or LAFW, but there was no significant difference between NOX2 shRNA animals and controls in any region. Taken together, NOX2 shRNA injection did not lead to significant changes in mRNA expression of ion channels thought to contribute to atrial ERP shortening in RAP.

## Discussion

In this study, we examine the role of oxidative injury in causing electrical remodeling in a RAP model of AF. We demonstrate that ROS generation in the atrium is a frequency-dependent phenomenon, with continued RAP leading to preferential elevation of mitochondrial ROS and NOX2 in the fibrillating atrium. To determine how oxidative injury may be creating a vulnerable substrate for AF, we examined which of the major ion channels invoked in causing ERP shortening in the RAP model (as well as in human AF) are sensitive to ROS inhibition. We discovered that the PKC_ɛ_-regulated, constitutively active *I*_KAch_ channel (*I*_KH_) was uniquely sensitive to inhibition of mitochondrial ROS and NOX2. To determine whether oxidative injury is involved in the initiation and/or maintenance of RAP-induced ERP shortening in the intact atrium – and resulting AF - we then performed targeted inhibition of ROS in the atria with a novel, gene-based approach. Specifically, targeted gene expression of NOX2 shRNA in the atria – to inhibit one of the major sources of ROS in the atria - led to a 113% increase in ERP when compared to control animals subjected to sustained duration of RAP. This prolongation of ERP was accompanied by an inability to induce persistent AF despite up to 12 weeks of continued RAP in animals that underwent atrial injection of NOX2 shRNA. See Figure 8C for a proposed model of the mechanisms by which NOX2 shRNA is attenuating electrical remodeling and consequent AF in the intact atrium.

### Likely sources of ROS that contribute to creation of a vulnerable AF substrate

ROS are unstable, reactive oxygen derivatives including O_2_^-^ and its derivatives hydrogen peroxide (H_2_O_2_), hydroxyl radicals (OH●), peroxinitrites (ONOO^-^) and hypochlorite ion (OCl^-^). They play significant roles in cardiac physiology as they act as crucial second messengers for growth and gene regulation (*25, 28, 29*). However, if the fragile balance of generation and neutralization of ROS is disturbed, excessive ROS can elicit pathologic cellular responses and lead to a number of cardiac diseases (*25, 28*).

Cellular, animal and clinical studies have shown high levels of ROS in the fibrillating atria, and suggested a role for oxidative injury in the pathophysiology of AF with stimulation of ROS sensitive kinase and phosphatases leading to electrical remodeling (*30–32*). In addition, oxidative injury may also promote adverse structural remodeling with inflammation ultimately leading to fibrosis, providing a substrate for AF persistence (*33, 34*). However, the precise role of oxidative injury in the genesis and maintenance of electrophysiological remodeling in AF and the specific enzymatic source(s) of oxidative injury involved herein have not been established (*18, 31, 35*). Whereas prior studies have established a role for ROS in modulation of several ion channels and EC coupling proteins, the data for ROS modulation of ion channels in the context of AF is limited (*20*). Finally, prior studies supporting a role for oxidative injury in the development or persistence of AF have been limited to the study of explanted atria or isolated cardiomyocytes; it remains unknown whether oxidative injury – by virtue of its effects on one or more atrial ion channels – affects electrical remodeling in the intact, fibrillating atrium. ROS generated in the cardiovascular system are primarily derived from NADPH oxidases (NOX), the mitochondrial electrical transport chain, xanthine oxidases and uncoupled eNOS (*15, 16, 36*). Among these enzymatic sources/complexes, NOX have emerged as a major initiating source for increased ROS generation in cardiovascular diseases. Moreover, mice with cardiac-specific overexpression of Rac-1, a necessary activator for NOX2, develop AF spontaneously (*17*), supporting a role for NOX2 in determining an atrial substrate for the new onset of AF. In humans, atrial NOX activity was independently associated with risk of developing AF after cardiac surgery (*37*), and atrium homogenates from AF patients contained more NOX-derived O2^-^ than controls (*9*). In this study, we confirmed a role for NOX2 in the initiation as well as the and maintenance of AF in the canine RAP model: attenuating NOX2 with an shRNA-based gene therapy approach not only delayed the onset of AF, but also prevented development of sustained AF. This effect appeared to be largely mediated by NOX2 shRNA preventing RAP induced ERP shortening. This suggests a role for NOX2 not only in post-operative AF, but also as a continuous trigger involved in the maintenance of persistent AF.

A further contribution of other ROS sources (mitochrondrial, xanthine oxidases, uncoupled eNOS) has also been hypothesized. Reilly et al. showed that while NOX2 was elevated in atria from patients with postoperative AF, uncoupled NOS and mitochondrial ROS were important sources of oxidative injury in patients with long standing AF (*18*). Indeed, our study also demonstrates a significant elevation of mitochondrial ROS in the rapidly paced atrium. Whether inhibition of mitochondrial ROS leads to beneficial effects on electrical remodeling similar to those noted in the current study with targeted inhibition of NOX2 remains to be determined. Even though our study did not show elevation of xanthine oxidase, Yoshizawa et al. recently demonstrated that the xanthine oxidase inhibitor febuxostat, in a short-term canine RAP study, prevented slowing in conduction velocity (paralleled by a reduction in interstitial fibrosis), but only had a minimal effect on ERP (*38*). This may point towards a differential effect of ROS on atrial electrophysiology based on cellular or subcellular origin. Alternatively, this may represent a pleiotropic effect of febuxostat.

### Ion channel(s) likely mediating ROS-induced ERP shortening in AF

In the absence of significant structural heart disease, electrical remodeling leading to AF is thought to be primarily driven by ERP shortening (*39–41*). We therefore chose to examine the role of OS in the genesis of AF in a canine RAP model of AF, in which ERP shortening is a major determinant of AF (*42–45*). The ERP is determined by the action potential duration (APD), which by itself is contingent upon the balance between inward currents (primarily Ca^2+^, which tends to keep the cell depolarized) and outward currents (primarily K^+^, which tends to repolarize) during the action potential plateau. RAP-induced APD shortening is multifactorial, involving both inward and outward currents. Multiple components of this fine balance have been hypothesized to participate in oxidative injury dependent ERP shortening, and we will review them below.

*I*_CaL_ is downregulated in persistent AF(*46*), an effect that we also demonstrated in this study (Supplementary Figure S1). Downregulation of *I*_CaL_ pore forming α-subunit mRNA appears to be the main contributor to this effect, but posttranscriptional mechanisms such as protein dephosphorylation and breakdown may also be involved (*26, 46, 47*). Carnes et al. demonstrated higher nitrosylation of the α1c subunit of L-type Ca^2+^ channels in the LAA of patients with long-standing AF(*48*). Nitrosylation appeared inversely related to cellular glutathione content, and superfusion of NAC resulted in an increase in *I*_CaL_. Notably, *I*_CaL_ did not appear to be sensitive to ROS inhibition in our model. Potential explanations for this discrepancy include species-related differences (canine versus human) or region of origin of atrial myocytes (left atrial appendage myocytes were studied by Carnes et al., whereas the current study examined posterior left atrial myocytes).

An increase in the inward rectifier current I_K1_ leading to a more negative resting potential is also described in AF. In particular, upregulation of Kir2.1 protein and mRNA has been described in humans with AF (*49–51*). Redox-dependent regulation of *I*_K1_ has been reported, with S-nitrosylation of the Cys76 residue of Kir2.1 increasing the channel open probability in isolated human atrial myocytes from patients in sinus rhythm (*52*). In our study, *I*_K1_ did not appear to be sensitive to ROS inhibition.

Among voltage gated K^+^ currents, the transient outward K^+^ current (*I*_to_) is consistently decreased in RAP due to reduction of K_v_4.3 expression (*45, 53*). Studies on ultra-rapid delayed rectifier K^+^ current (*I*_Kur_) point towards attenuation of this current in AF, though with some conflicting results (*54*). *I*_Kur_ was shown to be ROS-sensitive, with both S-nitrosylation and sulphenylation leading to reduced current density (*55, 56*). Overall, oxidation dependent regulation of K_v_1.5 which forms the ion conducting pore for *I*_Kur_, is unlikely to lead to ERP shortening (an opposite effect would be expected). Based on this rationale, we chose to forego evaluation of this channel in this study.

*I*_KACh_ causes APD shortening and cell membrane hyperpolarization. Increased vagal activity promotes AF by stabilizing atrial reentrant rotors (*57*). Patients with AF also display *I*_KH_ current (*22, 49, 58*). Whereas protein expression of Kir3 subunits is unchanged in RAP models (*58, 59*) and even decreased in AF patients (*50*), *I*_KH_ is increased due to enhanced open probability of the single channel secondary to slowed channel closure (*59*). Frequency-dependent membrane translocation of PKC_ɛ_ leads to abnormal channel phosphorylation that favors enhanced basal *I*_KH_ (*23*). In this study, we showed that *I*_KH_ was uniquely sensitive to ROS inhibition, as opposed to the acetylcholine-regulated fraction of *I*_KACh_. Furthermore, ROS inhibition prevented tachypacing induced membrane translocation of PKCε, providing a mechanistic basis for ROS-dependent *I*_KH_ activation. Since PKC isoforms – in particular PKC_ɛ_, - have been shown to not only be acutely sensitive to ROS (*24, 25*) but also lead to induction of ROS, especially in the setting of hypoxia (*60*) and in view of the fact that *I*_KACh_ protein subunits themselves were not noted to be oxidized in our study, our findings strongly support a mechanistic role for ROS in causing PKC_ɛ_ induced upregulation of *I*_KH_ in the fibrillating atrium.

### Oxidative injury –a potential therapeutic target in AF

Despite significant evidence that oxidative injury plays an important role in the generation of AF, clinical trials have provided only limited evidence for the benefit of conventional antioxidants. Carnes et al. treated a small group of patients undergoing cardiac surgery with ascorbic acid and demonstrated a 50% reduction of post-operative AF in the treated group when compared to controls (*61*). However, subsequent larger and randomized double-blind placebo-controlled trials with ascorbic acid produced mixed results (*35*). NAC also been used in several small clinical trials, but results have been mixed thus far (*35*). It has been argued that these studies were likely underpowered to see a significant effect. In addition, there is concern that systemically administered NAC does not reach a sufficient concentration at the myocardium for effective ROS scavenging. Instead of attempting to scavenge existing ROS, strategies aimed at preventing ROS formation might be more successful. For instance, statins reduce ROS formation by inhibiting Rac1 GTPase and hence decreasing NOX2 expression (*18, 62*). In small studies on patients undergoing cardiac surgery, there was an improvement in redox state after statin therapy. A meta-analysis showed a significant association between statin use and reduction in post-operative AF and secondary prevention of AF, but no difference in primary prevention of AF (*63*). A subsequent blinded, placebo controlled randomized trial did not reveal any effect of statin therapy on post-operative AF (*64*). In addition to statins, the renin-angiotensin-aldosterone system has also been proposed as an upstream regulator of NOX, and hence a potential target for OS-induced AF (*31*). Several meta-analyses have shown an overall beneficial effect of RAS inhibition on incidence or recurrence of AF, though with significant heterogeneity (*65, 66*). However, it remains difficult to separate the antioxidant effect of RAS inhibition with its multiple other effects on the cardiovascular system.

This variable efficacy of small molecule approaches is likely at least in part secondary to variable drug bioavailability and distribution, affinity for the target, and pleiotropic effects. As opposed to pharmacological therapy, gene therapy has the advantage of a localized, upstream effect allowing for targeted suppression of one or more major enzymatic sources of oxidative injury. The present study suggests efficacy of NOX2 shRNA gene transfer for treatment of sustained AF, with a durable effect lasting at least 12 weeks in the canine RAP model. In the next section we discuss the potential translational utility of such an approach in patients with AF.

### A targeted, gene based therapy for AF – is it time?

Gene therapy for AF has been investigated over the last decade, but has remained in pre-clinical testing. ERP prolongation was targeted by gene therapy with an adenovirus vector expressing a dominant-negative mutant of the *I*_Kr_ channel KCNH2. Gene vector was delivered in a pig model of RAP-induced AF either with the gene painting method (*67*) or with direct myocardial injection followed by electroporation (*68*). Both groups showed prolongation of the action potential duration or ERP. Short-term expression of the vector (<3 weeks) was present in both studies, with a delay in AF induction by 5-10 days. Conduction velocity was also targeted by using an adenovirus vector expressing atrial connexins, either Cx40 (*69*) or Cx43 (*69, 70*) in pig models of RAP-induced AF. Both studies showed improved conduction velocity with gene therapy, and short-term efficacy with reduced AF induction up to 7 days (*69*) or 14 days (*70*). Other mechanisms such as cardiomyocyte apoptosis were also targeted with a similar approach: an adenovirus vector expressing a silencing RNA (siRNA) for Caspase 3 administered via myocardial injection and electroporation in a pig model of RAP-induced AF. This resulted in improved conduction velocity and short-term reduction in AF inducibility (14 day follow-up). Vagal signaling was also targeted by using a non-viral vector expressing C-terminal peptides of Gα subunits in a canine model (*71*). This short-term study (3 days) showed effective attenuation of vagal-induced ERP shortening and AF inducibility. Finally, atrial fibrosis was targeted in the canine rapid ventricular pacing model of heart failure by injection and electroporation of a dominant negative TGF-β type II receptor (*72*). This study showed a decrease in interstitial fibrosis associated with reduction in conduction inhomogeneity and decreased duration of AF after 3-4 weeks of rapid ventricular pacing.

While promising, previously published gene therapy for AF has been mostly focused on short-term efficacy. In this study, we show that NOX2 shRNA has a long-lasting effect of at least 12 weeks after initiation of RAP with a marked and sustained reduction in AF burden.

Demonstration of longer term gene expression with our non-viral approach is therefore an important step towards translation to humans with AF. While viral vectors – specifically adeno-associated virus and lentivirus – have been shown to have long lasting gene expression (*73*), a non-viral approach may have certain advantages compared to viral approaches e.g. reduced inflammation, less probability of gene spillover to surrounding myocardium, etc. Nonetheless, the need for electroporation to facilitate non-viral gene delivery does add to the complexity of this approach, and requires the development of surgical or per-cutaneous devices that can facilitate safe and effective electroporation in the human atrium. Percutaneous access to the pericardial space is now routinely used for numerous minimally invasive surgical applications and arrhythmia ablations. Adapting the existing tools for minimally invasive surgery to the gene transfer application should allow a broader application of this technique.

Since NOX2 induced oxidative injury has been shown to induce pro-fibrotic signaling in the heart (*74, 75*), targeting NOX2 with a gene-based approach would also be expected to reduce fibrosis in the AF atrium. Since fibrosis is not a key feature of the RAP model, we did not examine the effect of NOX2 shRNA on atrial fibrosis in this model (see Limitations).

### Limitations

Due to the nature of sub-epicardial gene injection, it is conceivable that target expression is somewhat heterogeneous. Whereas it is difficult to examine the distribution of transcribed gene product with an shRNA-based approach, we did demonstrate fairly homogeneous suppression of oxidative injury in regions injected with NOX2 shRNA. In addition, the approach used in the current study allowed at least moderately homogeneous atrial gene expression in prior studies (*71, 72*). Future studies are warranted to more systematically examine the degree of gene expression homogeneity required for a beneficial therapeutic effect.

AF maintenance is promoted by ERP shortening, but also by slowing and inhomogeneity of conduction. As described earlier, strategies aimed at improving conduction velocity (e.g. Cx40 or Cx43 gene transfer) were able to delay or prevent AF induction. We did not evaluate the effect of NOX2 shRNA on conduction velocity, and therefore cannot exclude an additional mechanism leading to reduced AF burden in this model.

Since the RAP model is primarily a model of electrical remodeling in AF, we did not examine the effect of NOX2 shRNA on atrial fibrosis. Future studies need to examine the effect of this gene therapy approach on atrial fibrosis in a more relevant model such as the canine heart failure model of AF.

## Materials and Methods

### Study design

The objective of this study is to determine the precise role of oxidative injury in causing electrical remodeling in the intact, fibrillating atrium. Our pre-specified hypotheses were: 1) Upregulation of *I*_KH_ – an ion channel thought to be an important contributor to electrical remodeling in AF –is mediated by a frequency-dependent increase in oxidative injury in the setting of AF, with resulting activation of PKC_ɛ_ and 2) oxidative injury leads to not only the initiation but also the maintenance of ERP shortening in the intact, fibrillating atrium.

This was an experimental study that was performed in large animals, specifically dogs and pigs. Dogs and pigs used in this study were maintained in accordance to the Guide for the Care and Use of Laboratory Animals published by the U.S. National Institutes of Health (NIH Publication No. 85-23, revised 1996) as approved by the IACUC of the Northwestern University.

A total of 32 dogs were used for this study. Dogs used were either normal controls (N = 8) or were subjected to RAP (N = 24). In control animals, the following types of experiments were performed: i) tissue analysis (n = 5), ii) atrial myocyte isolation (n = 6). Atrial myocytes isolated from normal dogs were subjected to *in vitro* tachypacing or to single cell electrophysiological analysis. Participation of PKC_ε_ in the induction of *I*_KH_ by oxidative injury was assessed in *in vitro* tachypaced atrial myocytes.

Dogs were subjected to rapid atrial pacing (RAP) for a period of 3-12 weeks. A subset of dogs undergoing RAP also underwent gene injection. The RAP dogs were divided into three groups: Group 1 - No gene injection (n = 15); Group 2 - NOX2 shRNA injection (n = 6); and Group 3 - Scrambled gene injection (n = 3). After pacemaker implantation, all RAP dogs underwent periodic pacemaker interrogations to assess for duration of induced AF. In dogs that underwent gene injection, NOX2 shRNA or scrambled shRNA was injected sub-epicardially in canine atria prior to initiation of RAP. NOX2 shRNA was used to inhibit NOX2, a major enzymatic source of oxidative injury in the AF atrium. One week after gene injection, RAP was initiated and continued for 3-12 weeks to assess for development of AF. At the time of terminal surgery, each group of RAP dogs underwent one or more of the following procedures (the procedures are not mutually exclusive): Group 1: Electrophysiological testing (includes ERP analysis, AF recordings) (n = 11); Tissue analysis (n=8); Atrial myocyte isolation (n = 8).

Group 2: Electrophysiological testing (includes ERP analysis, AF recordings) (n = 4); Tissue analysis (n=5). Group 3: Electrophysiological testing (includes ERP analysis, AF recordings) (n = 1); Tissue analysis (n=1).

Atrial myocytes isolated from RAP dogs were subjected to single cell electrophysiological analysis. Sensitivity of specific ion channels to ROS inhibition was determined by whole cell patch clamp experiments in control and RAP atrial myocytes, performed in the presence of different ROS inhibitors. Lastly, atrial myocytes were isolated from one pig and subjected to increasing frequency of *in-vitro* tachypacing, to assess for ROS generation.

See Supplementary Materials for detailed Methods.

### Statistical Analysis

All data is presented as mean ± SEM.

For each ion current, current amplitude was compared at each voltage step by unpaired t-tests.

To assess differences in ERP between different groups of animals, individual ERPs obtained in each animal were combined by region and group, to determine a statistically significant difference in means between treatment groups and between atrial regions by using two-way ANOVA with Holm-Sidak method for pairwise comparisons.

Time to sustained AF between the control and active gene groups was compared by log-rank tests on interval censored data. Time to AF was compared for both short duration AF (>30 minutes) and long duration AF (>8 hours). Due to censoring of observations, the median time in AF per occasion per dog was calculated. Difference between groups was determined by a Wilcoxon rank sum test.

AF characteristics were compared between treatment groups and between atrial regions (PLA, LAA) by using two-way ANOVA with Holm-Sidak method for pairwise comparisons. Mean fluorescence in isolated atrial myocytes was compared using one way ANOVA. Other cellular and tissue parameters (O^2-^ levels, density of protein bands on western blot, mRNA levels by PCR, 8-oxo-DG stained nuclei on immunohistochemistry) were compared between treatment groups by unpaired t-tests.

Gene and protein expression were compared by unpaired t-tests or ANOVA with Holm-Sidak method for pairwise comparisons when more than two groups were examined. The values were considered significantly different at p < 0.05.

## Supplementary Materials

### Materials and Methods

Fig. S1. No effect of mitochondrial ROS inhibition on *I*_CaL_ in RAP myocytes.

Fig. S2. PKC mediated attenuation of *I*_KH_ is at least partially ROS mediated.

Fig. S3. NOX2 shRNA and scrambled shRNA constructs and *in vitro* testing.

Fig. S4. Ion channels mediating ERP shortening are not attenuated by NOX2 shRNA gene injection *in vivo*.

Table S1. Echocardiographic evaluation of NOX2 shRNA and control animals.

Table S2. qRT-PCR primer sequences.

Supplemental References: 76-79

## Author contributions

Conceptualization, SY, AP, GLA, RA.; Methodology, SY, AP, JH, JN, JAW, GLA, RA Investigation, SY, AP, JH, WZ, JN, AB, DAJ, GG, TW, SB, BB, RA.; Writing – Original Draft, SY, AP, GLA, RA.; Writing – Review & Editing, SY, AP, JAW, GLA, RA, BPK, RP; Funding Acquisition, AP, RA; Supervision, GLA, RA.

