## Supplemental Material for "Disruption of NOX2-dependent Oxidative Injury with a Targeted Gene-Therapy Approach Prevents Atrial Fibrillation in a Canine Model"

This file includes:

### Materials and Methods

*Animal experimentation*

Retired breeder female hound dogs (age 1-4 years, weight 22-33 kg) and a swine (weight 43 kg) used in this study were maintained in accordance to the Guide for the Care and Use of Laboratory Animals published by the U.S. National Institutes of Health (NIH Publication No. 85-23, revised 1996) as approved by the IACUC of the Northwestern University. Before undergoing the procedures listed below, all animals were premedicated with acepromazine (0.01 – 0.02 mg/kg) and were induced with propofol (3-7 mg/kg). All experiments were performed under general anesthesia (inhaled) with isoflurane (1-3 %). Adequacy of anesthesia was assessed by toe pinch and palpebral reflex.

*Assessment of superoxide generation*

Frozen tissue samples were crushed and rotor homogenized with protease inhibitor (Halt protease and phosphatase inhibitor cocktail, Thermo-Scientific). Protein concentration was determined using Pierce BCA Protein Assay Kit (Thermo-Scientific). Lucigenin (5 μmol/L, Enzo Life Sciences) and NADPH (100 μmol/L, Calbiochem) were each added in the presence and absence of the following inhibitors: apocynin (NOX2), mito-TEMPO (mitochondrial reactive oxygen species (ROS) scavenger), L-NMMA (nitric oxide synthase) and oxypurinol (xanthine oxidase). The photon outputs were measured using a luminometer (Berthold Technologies, LUMAT LB 9507) (*76*).

*Atrial Cardiomyocyte Isolation*

While the dog was still deeply anesthetized, the hearts was quickly removed and immersed in cold cardioplegia solution containing (mM) NaCl 128, KCl 15, HEPES 10, $MgSO_4$ 1.2, $NaH_2PO_4$ 0.6, $CaCl_2$ 1, glucose 10, and heparin (0.001 U/mL); pH 7.4. All solutions were equilibrated with 100% $O_2$. The aorta was cannulated, and the heart was perfused with cold cardioplegia solution until effluent was clear of blood and heart was cold (5-10 min). The ventricles were cut away, the left circumflex coronary artery was cannulated, and the left atrium and right atrium were dissected free. The atria were slowly perfused with cold cardioplegia while leaks from arterial branches were ligated with suture to assure adequate perfusion. The atria were then perfused with Tyrode's at 37 °C for 5 min to remove cardioplegia solution and assess for viability—i.e., the reestablishment of beating. If viable, the atria were then perfused at ~12 mL/min with $Ca^{2+}$-free Tyrode's solution for ~20 min, followed by ~40 min of perfusion with the same solution containing Liberase (Liberase TH Research Grade, Roche 05401151001) and 1% BSA; all at 37°C. Thereafter, the atrial tissue was transferred to dish and cut into small pieces (~0.5 $cm^2$). These tissue pieces were then transferred to conical plastic tubes, and fresh enzyme solution (37 °C) was added. The tissue pieces were triturated in the fresh enzyme solution for 5-15 min. The triturated tissue suspension was then filtered through nylon mesh (800 μm). The filtered cell tissue suspension was briefly centrifuged at ~500 g, then enzyme solution poured off, and cell tissue suspension resuspended in Tyrode's solution containing 250 μM $Ca^{2+}$ and 0.1 % BSA. This resuspension was then filtered through a nylon mesh (210 μm), centrifuged at 500 g, and again resuspended in Tyrode's solution containing 250 μM $Ca^{2+}$ and 0.1% BSA to isolate dispersed cells. After cells settled for about 30 minutes, the solution was suctioned off and gradually replaced with a HEPES-buffered solution containing (mM) NaCl 137, KCl 5.4,

$MgCl_2$ 1.0, $CaCl_2$ 1.8, HEPES 10, glucose 11, and 0.1% BSA; pH 7.4. After isolating the cardiomyocytes, $Ca^{2+}$ concentration was raised to 1.8 mM in 1X Tyrodes Solution.

*Single cell electrophysiology*

Whole cell patch clamp experiments were conducted using Axon 200A amplifier interfaced to personal computer equipped with Digidata 1322A and pClamp 9 software (Axon Instruments). At 37◦C, recordings were made from single, quiescent, and rod-shaped myocytes. Recording electrodes were fabricated from borosilicate glass with 1.5-3.0 MΩ resistance. Pipette solution for inwardly rectifying currents contained (mM: K-Asp, 110; KCl, 20; $MgCl_2$, 1.04; Mg-ATP, 5; Lt-GTP, 0.1; HEPES, 10; Na-phosphocreatine, 5; EGTA 5; pH 7.3 with KOH). Signals were low-passed filtered at 1 kHz and digitized at 10 kHz and current recordings were not corrected for leak. Cell capacitance was evaluated by nullifying the area under the capacitive transient elicited by a -10 mV test pulse. External recording solution for inwardly rectifying currents contained (mM: NaCl, 136; KCl, 5.4; $MgCl_2$, 1.0; NaHPO4, 0.33; HEPES, 5.0; $CaCl_2$, 1.0; glucose, 10; 4AP, 2.0; $CdCl_2$, 0.2; pH 7.35 with NaOH). Inwardly rectifying currents were recorded by applying step pulse protocol from -120 mV through -20mV for 4 second from a holding potential of -40 mV or 400 ms ramp pulse protocol from a holding potential of -40 mV to voltage between 10 mV to -120 mV. All the currents are expressed as densities (pA/pF). $I_{KH}$ and $I_{KACh}$ were measured as the 100 nM TQ- and 10 μM CCh- sensitive current, respectively by calculating differences between the currents before and after the application of drug. $I_{K1}$ was referred as residual inward current after TQ. Following drugs are applied: BIM1 (Cayman chemical), gp91-tat (Anaspec) and $PKC_\varepsilon$ inhibitory peptide (H-Glu-Ala-Val-Ser-Leu-Lys-Pro-Thr-OH, Santacruz).

For L-type $Ca^{2+}$ currents measurements, pipette solutions contained (mM: Cs-Asp, 120; CsCl, 25; $MgCl_2$, 1.5; NaCl, 6; K2-ATP, 4; EGTA, 0.056; HEPES, 20; pH 7.2 with CsOH). L-type $Ca^{2+}$ currents were measured as a 10 μM nifedipine sensitive currents by applying 250 ms ramp pulse protocol from a holding potential of -80 mV to voltage between 20 mV and -120 mV. External solution for L-type $Ca^{2+}$ currents measurements contained (mM: NaCl, 140; KCl, 5.4; $MgCl_2$, 0.5; HEPES, 10; $NaH_2PO_4$, 0.4; 4-AP, 4; Glucose, 11; $CaCl_2$, 1.8; $BaCl_2$, 0.25; pH 7.4 with NaOH).

*Primary culture and In vitro-Tachypacing*

Freshly isolated atrial cardiomyocytes from control dogs and control pigs were plated onto laminin-coated (1 μg/$cm^2$) multiwell plates with M199 plating media (Invitrogen) supplemented with 10 % FBS, 50 IU/mL penicillin, 50 μg/mL streptomycin, and ITS (Insulin-transferring-Sodium Selenite, Sigma) in mammalian tissue culture incubator at 37 ◦C, 95% $O_2$/5% $CO_2$.

For PKCε membrane translocation experiments, after 1hr incubation, dead and unattached canine cardiomyocytes were removed and fresh media was added. Cells were then paced at either 1 Hz or 3 Hz for 6-12 hrs using square wave 5 ms pulses driven via IonOptics C-PACE stimulator. For some experiments, 10 mM NAC was added in the media. After in-vitro tachypacing, cells were snap-frozen for subsequent immunoblot analysis.

For the ROS measurement experiment, isolated pig atrial myocytes were then incubated with a ROS-sensitive dye, 5 μM CellROX Deep Red (Thermo Scientific) and subjected to: a) no pacing, b) *in-vitro* pacing at 1 Hz and c) *in-vitro* pacing at 2 Hz for 6 hours. Fluorescent signal was measured using confocal microscope (Zeiss 510 Meta).

*Immunoblots*

Total protein extracts were extracted from snap-frozen canine atria by using the lysis buffer containing 20 mM Tris-HCl (pH 8.0), 100 mM NaCl, 1 mM EDTA, 0.5 % NP-40 and 1 x protease inhibitor cocktail solution. Protein concentration was determined using Pierce BCA Protein Assay Kit (Thermo-Scientific) and BSA was used as a standard. Total protein extracts of 20-30 µg were separated on 10 % SDS-PAGE gels, and transferred to polyvinylidene difluoride membranes (Immun-Blot PVDF Membrane, BioRad). The membranes were incubated with 5 % nonfat dry milk in PBST (phosphate-buffered saline (PBS), 0.05 % (v/v) Tween-20 (Sigma), pH 7.4), and then probed with the primary antibody and horseradish peroxidase-conjugated secondary antibodies. The protein signal was visualized by using the ECL detection system (Amersham Biosciences). The membranes were re-probed using anti-cadherin, GAPDH or HSP90 antibodies, which serve as loading control. All results were scanned and quantified by ImageJ.

*Preparation of cytosolic and membrane fractions*

We used Thermo Mem-PER plus membrane protein extraction kit (Thermo Scientific) for separating the membrane fraction from the cytosolic fraction through phase partitioning. Isolated- and differentially paced- atrial cardiomyocytes or whole tissue lysates were resuspended in cell wash solution and centrifuged at 300 g for 5 min. The supernant was discarded and permeabilization buffer was added to the pellet. Permeabilized samples were centrifuged for 15 min at 16000 g. The supernatant containing cytosolic proteins was saved for later use. Solubilization buffer was added to the pellet and resuspended mixture was incubated at 4 C for 30 min and then centrifuged at 16,000 g for 15 min at 4 C. Supernatant containing solubilized membrane protein was saved. The cytosolic and membrane fractions were used for immunoblot analysis with anti-$PKC_\varepsilon$ antibody as described above.

For $PKC_\varepsilon$ membrane localization experiments, the intensity of the cytosolic and membrane fraction immunoreactive bands were determined and normalized to their respective loading control. Membrane fraction is expressed as percentage of total content (membrane + cytosol fractions).

*Cryosectioning and Immunohistochemistry*

Canine atrial tissue was excised and PLA, LAFW, LAA, posterior right atrium (PRA), right atrial free wall (RAFW) and right atrial appendage (RAA) regions were dissected. The preparations were frozen in OCT tissue freezing medium (VWR) at ~-50 ºC in 2-methyl butane cooled by dry ice, and stored at -80°C until use. The frozen preparations were secured on the chuck of a cryostat with tissue-freezing medium and serially sectioned (at - 25 °C) at 10 µm thickness. Sections were mounted on Superfrost Plus slides (VWR) and stored at - 80 °C until use.

Sections taken from -80 ºC freezer were air-dried and underwent fixation with 75 % acetone/ 25 % ethanol and washed 3 times in TBS-T. The sections were then treated in 3% hydrogen peroxide. After washing three times in TBS-T, the sections were blocked in protein block reagent (Dako) and then incubated with primary antibodies diluted in antibody diluent reagent (Dako) in a humid box at -4 °C overnight. The sections were washed three times in TBS-T, and incubated with Dako envision secondary antibodies in a humid box at RT in the dark for 30 min. After washing three times in TBS-T, the sections were dehydrated with series of Ethanol and Xylene and mounted with cytoseal (VWR). Stained sections were visualised using transmitted light microscope (Olympus) or TissueFax system (TissueGnostics). Aquired images were analyzed by histoquest software (TissueGnostics).

*Plasmid preparation*

The plasmids pLKO-shNox2, encoding NOX2 shRNA (Dharmacon # RHS3979) and pLKO-shSC a nontargeting control (Dharmacon # RHS6848) were confirmed by sequencing (see Supplementary Figure S3 for shRNA sequence). Plasmids were transfected with lipofectamin (Invitrogen) into HEK293 cells cultured in DMEM media. Efficiency of NOX2 knockdown was confirmed by qRT-PCR. Large scale of plasmids were prepared using the Qiagen Endo-free Giga kit (Qiagen 12391) with purity of DNA OD260/OD280~1.8.

*Quantitative real-time PCR*

Reverse transcription and quantitative RT-PCR (qRT-PCR) were performed using SYBR Green reagents (Invitrogen). Primers for all gene expression experiments are listed in Table S2. Samples were analyzed on an ABI 7500 sequence detection system (Applied Biosystems). Abundance of mRNA was normalized to TBP (TATA boxing binding protein). For NOX2, internal LAA was used as comparator. For all other genes, control LAA was used as comparator.

*Echocardiography*

Comprehensive echocardiography was performed prior to pacemaker implantation and immediately prior to the terminal study. Echocardiographic data included left ventricular end-diastolic and systolic dimensions, left ventricular ejection fraction, left ventricular global longitudinal strain, right ventricular size and function, left atrial volume and left atrial strain.

*Pacemaker implantation*

The right jugular vein was accessed by direct cutdown and ligated distally. A bipolar screw-in Medtronic pacing lead was inserted through an incision in the right jugular vein. The tip of the lead was fluoroscopically placed and fixed in the RA appendage after confirming adequate capture threshold (<0.5 mV with pulse width 0.4 ms). The proximal end of the pacing lead was connected to a custom-modified Medtronic programmable pulse generator that was subsequently implanted in a subcutaneous pocket in the neck. After all the incisions were closed, the dogs were allowed to recover from anesthesia and were returned to the animal quarters.

*Gene injection*

A lateral thoracotomy was performed under general anesthesia as detailed above. NOX2 shRNA or scrambled shRNA (5-7.5mg per chamber) was injected in the atria. For the first 4 animals, gene injection was limited to the PLA. Subsequent animals received NOX2 shRNA in the entire left and right atria, omitting only the right atrial appendage to avoid interference with the atrial pacing lead. ShRNA was diluted in sterile saline for a final concentration 0.6-0.9 mg/mL. A volume of approximately 0.5 mL was injected subepicardially at multiple sites spaced 0.5-1cm apart. Immediately after gene injection, electroporation was performed at each site of injection as follows: Two gold-plated, needle-style electrodes (10-mm length each) or a Coolrail Linear Pen (Atricure) were placed at each gene injection site. Electroporation was performed as previously described with eight pulses of 1 second at 120–150 V/cm2 (ECM 830, Harvard Bioscience). After gene delivery, the chest was closed and the animal was allowed to recover.

*Rapid atrial pacing*

One week after recovery from gene injection, rapid atrial pacing was initiated. After confirming adequate threshold for atrial capture (<0.5 mV with pulse width 0.4 ms), rapid atrial pacing was performed incrementally over 1-3 days until adequate capture was confirmed at 600 bpm. Pacing was interrupted three times weekly for up to 30 minutes to quantify AF duration over a period of 3-4 weeks (short term experiments) or once weekly for up to 8 hours for 12 weeks (long-term experiments).

*Terminal electrophysiological study*

A lateral sternotomy was performed under general anesthesia as detailed above. A terminal EP study was performed and all data were acquired by a 128-channel mapping system (PruckaCardioLab) at a sampling rate of 977 Hz. If sinus rhythm was present at baseline, effective refractory period (ERP) was determined. If the animal was in AF, attempt was made to terminate AF with burst pacing or DCCV. ERPs were determined in the PLA and LAA, at a baseline cycle length of 400 msec; starting at 10 msec, an extrastimulus (S2) was delivered at 10 msec increments, until atrial capture was obtained. Periods of AF lasting more than 10 seconds were recorded and analyzed as described in the section on AF electrogram analysis.

Upon finishing the *in vivo* portion of the study, and after confirming a very deep plane of anesthesia, the heart was removed and perfused with cold cardioplegia solution. The atria were dissected, snap frozen, and subjected to further analysis as detailed in the previous methods sections.

AF Electrogram Analysis:

AF episodes lasting more than 10 seconds were recorded in order to determine the following electrogram characteristics: 1) Dominant Frequency (DF), 2) Organization Index (OI), 3) Fractionation Interval (FI), 4) Shannon's Entropy (ShEn) and 5) Recurrence Cycle Length ($CL_R$). Briefly, DF is a frequency domain measure of activation rate. OI is a frequency domain measure of temporal organization or regularity. FI is the mean interval between deflections detected in the electrogram segment. ShEn is a statistical measure of complexity. $CL_R$ is the cycle length of the most recurrent morphology.

**Dominant Frequency (DF).** DF is a frequency domain measure of activation rate. Following bandpass filtering with cutoff frequencies of 40 and 250 Hz and rectification, the power spectrum of the electrogram segment was computed using the fast Fourier transform. The frequency with the highest power in the power spectrum was considered the DF.

**Organization Index (OI)**. OI is a frequency domain measure of temporal organization or regularity (*77, 78*). It has been shown that AF episodes with recordings with high OI are more easily terminated with burst pacing and defibrillation. OI was calculated as the area under 1-Hz windows of the DF peak and the next three harmonic peaks divided be the total area of the spectrum from 3 Hz up to the fifth harmonic peak.

**Fractionation Interval (FI).** FI is the mean interval between deflections detected in the electrogram segment. Deflections were detected if they met the following conditions: 1) the peak-to-peak amplitude was greater than a user determined noise level, 2) the positive peak was within 10 ms of the negative peak, and 3) the deflection was not within 50 ms of another deflection. The noise level was determined by selecting the amplitude level that would avoid detection of noise-related deflections in the iso-electric portions of the signal. FI is dependent on both the AF cycle length and the fractionation of the electrogram (*79*).

**Shannon's Entropy (ShEn).** ShEn is a statistical measure of complexity (*79*). The 4000 or 3908 (depending on the 1kHz or 977 Hz sample rate) amplitude values of each EGM segment were binned into one of 29 bins with width of 0.125 standard deviations.

ShEn was then calculated as:

$$ShEn = \frac{-\sum_{i=1}^{29} p_i \log_{10} p_i}{\log_{10} p_i}$$

In this equation, pi is the probability of an amplitude value occurring in bin *i*.

*Antibodies*

Primary antibodies: rabbit polyclonal anti $PKC_{\varepsilon}$ (cell signaling 2683), mouse monoclonal anti 8-OHdG (JalCA, MOG-020P), rabbit polyclonal anti-NOX2 (gp-91, Santa Cruz sc-20782), rabbit monoclonal anti-HSP90 (Cell Signaling 4877), rabbit monoclonal anti-GAPDH (cell signaling 5174), mouse monoclonal anti-pan Cadherin (Abcam ab6528). Secondary antibodies: anti-mouse IgG conjugated HRP (Jackson Immunoresearch), anti-rabbit IgG conjugated HRP (Jackson Immunoresearch).

**Supplementary Figures**

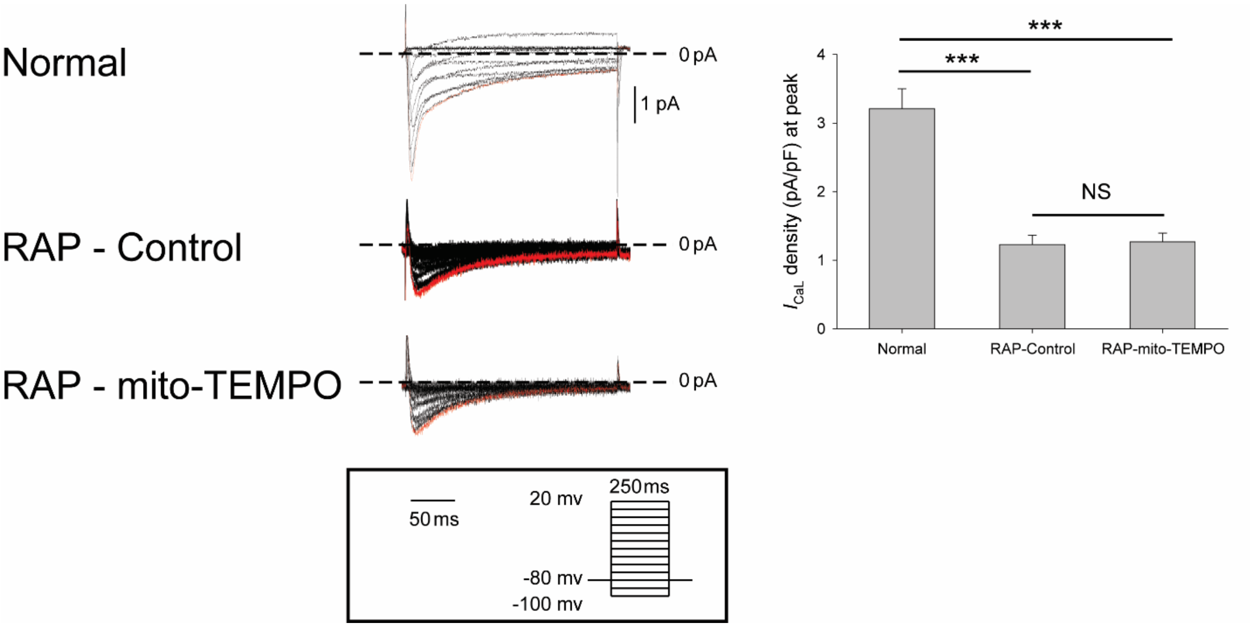


**Fig. S1. No effect of mitochondrial ROS inhibition on $I_{CaL}$ in RAP myocytes**. Raw traces (left panels) of $I_{CaL}$ elicited by 250 ms step pulses from a holding potential of -80 mV to voltage between -100 mV and 20 mV (pulse protocol shown in inset) and the current density at peak (right panels) for normal, control RAP and mito-TEMPO preincubated RAP atrial myocytes.

Data are presented as mean ± SEM; *** $p < 0.001$.

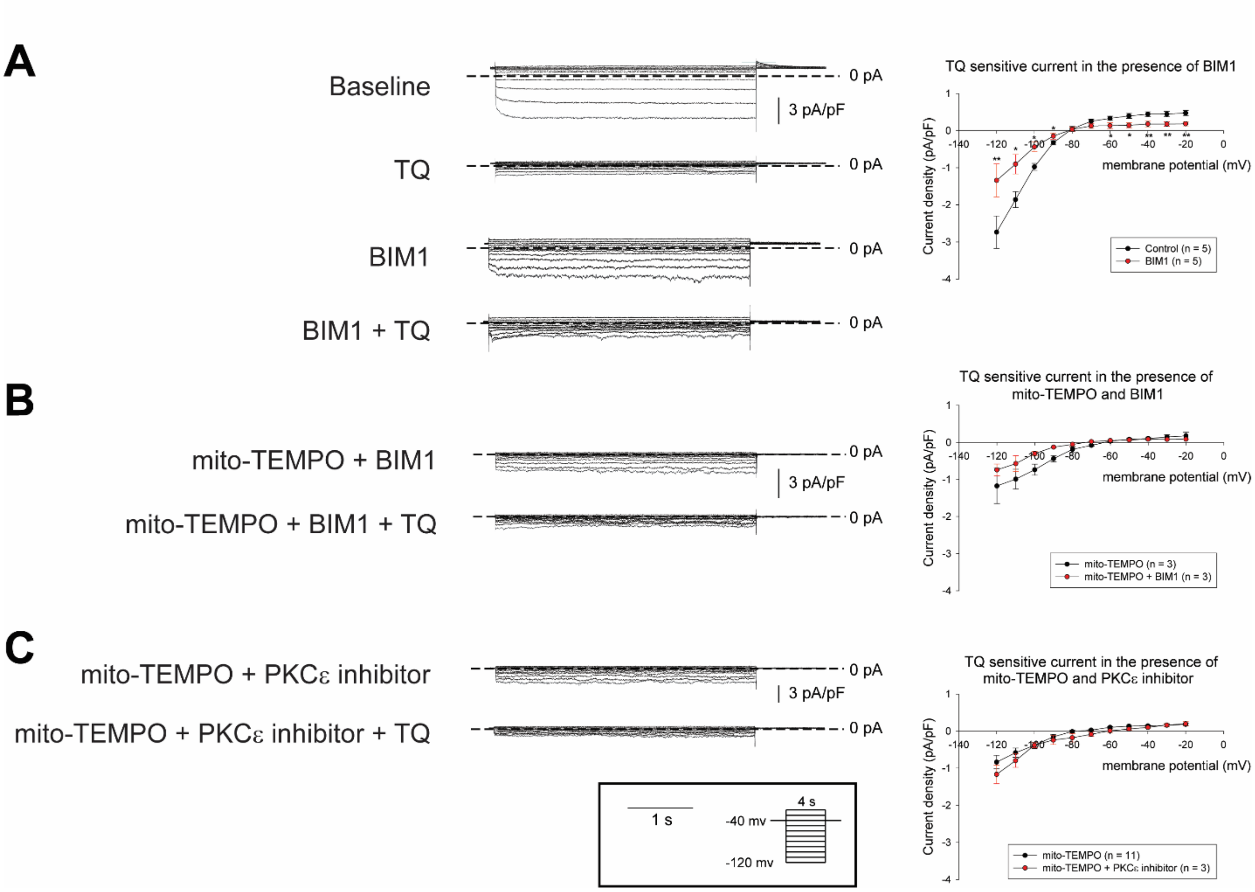


**Fig. S2. PKC mediated attenuation of $I_{KH}$ is at least partially ROS mediated.** (A - C) Raw traces (left panels) of $I_{KH}$ in the presence of (A) BIM1, (B) BIM1 + mito-TEMPO and (C) PKCε inhibitor + mito-TEMPO elicited by 4 seconds step pulses from a holding potential of -40 mV to voltage between -120 mV and -20 mV (pulse protocol shown in inset) and I-V curve (right panels). Data in I-V plots are presented as mean ± SEM at given membrane potentials; * $p < 0.05$ and ** $p < 0.01$.

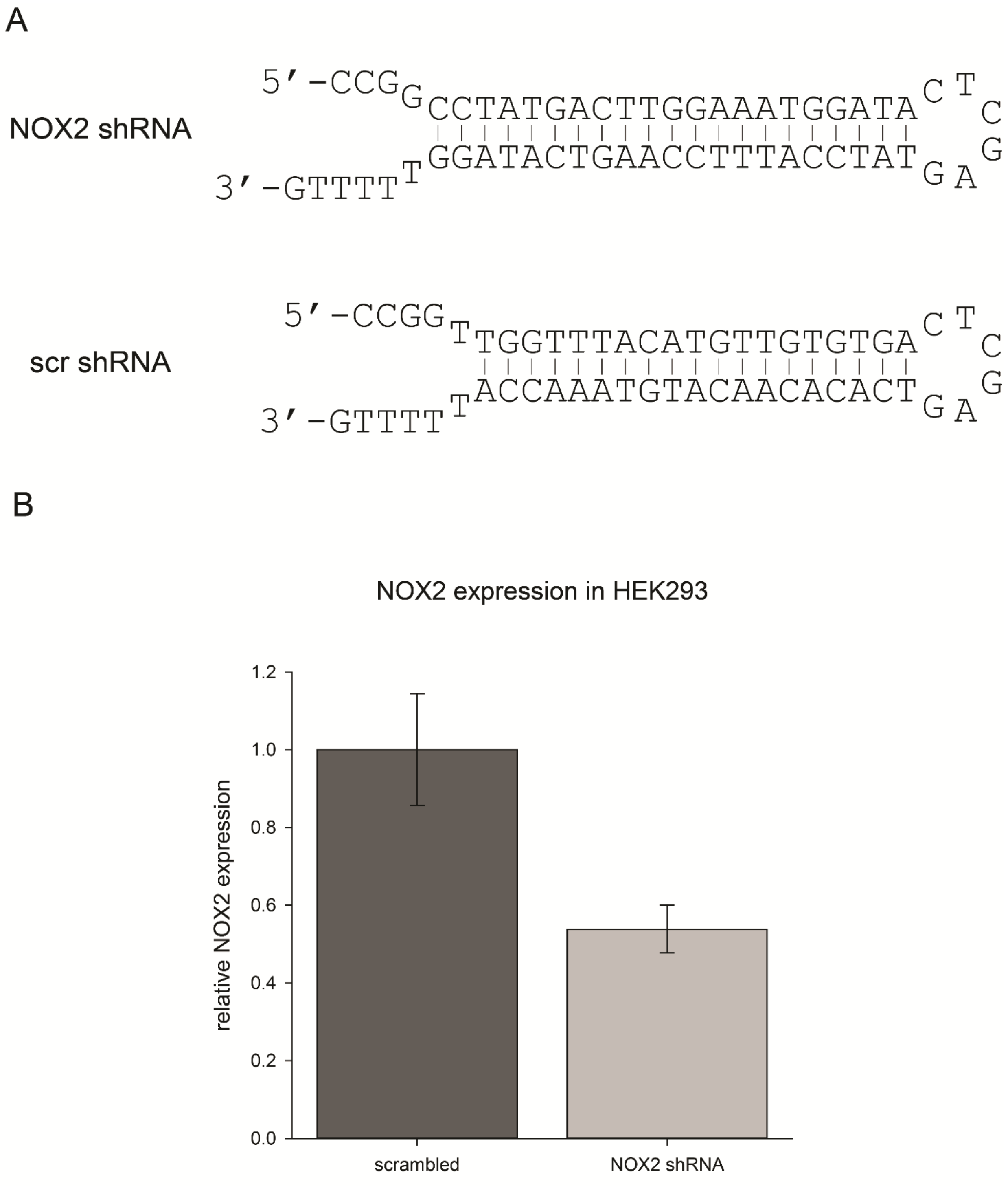


**Fig. S3. NOX2 shRNA and scrambled shRNA constructs and *in vitro* testing.** (A) Illustration of the NOX2 shRNA and scrambled shRNA constructs. (B) NOX2 expression was examined by qPCR in control and NOX2 shRNA-transfected HEK293 cells. N=2.

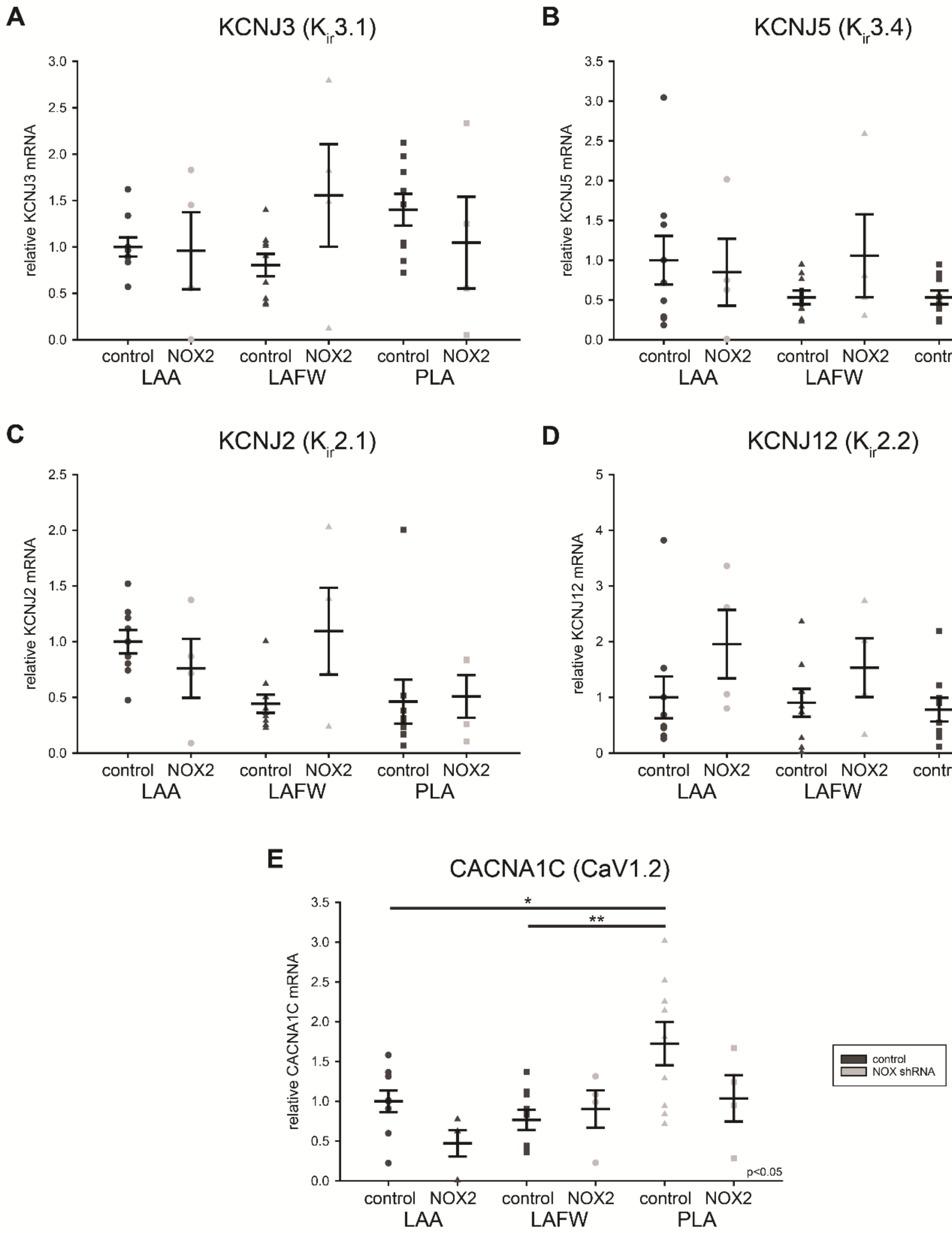

A
KCNJ3 ($K_{ir}$3.1)
relative KCNJ3 mRNA
B
KCNJ5 ($K_{ir}$3.4)
relative KCNJ5 mRNA
C
KCNJ2 ($K_{ir}$2.1)
relative KCNJ2 mRNA
D
KCNJ12 ($K_{ir}$2.2)
relative KCNJ12 mRNA
p<0.05
E
CACNA1C (CaV1.2)
relative CACNA1C mRNA
*
**
p<0.05
control NOX2 control NOX2 control NOX2
LAA LAFW PLA
control
NOX shRNA

**Fig. S4. Ion channels mediating ERP shortening are not attenuated by NOX2 shRNA gene injection *in vivo*.** (A) KCNJ3, (B) KCNJ4, (C) KCNJ2, (D) KCNJ12 and (E) CACNA1C expression was examined by qPCR in left atrial regions (LAA: left atrial appendage, LAFW: left atrial free wall, PLA: posterior left atrium) of control (n = 9) and NOX2 shRNA (n = 4) animals. Data are mean ± SEM; * $p < 0.05$ and ** $p < 0.01$. ANOVA significance shown in plot.

**Supplementary tables**

**Table S1.** Echocardiographic evaluation of NOX2 shRNA and control animals.

| | Control (n=5) | | | NOX2 shRNA (n=3) | | | | |
|---|---|---|---|---|---|---|---|---|
| | baseline | terminal | p Vs baseline | baseline | p Vs control | terminal | p Vs baseline | p Vs control |
| LA min volume (mL) | 13.2 ± 4.3 | 27.0 ± 18.4 | 0.16 | 9.7 ± 2.1 | 0.29 | 18.3 ± 6.3 | 0.17 | 0.52 |
| LA max volume (mL) | 23.2 ± 6.4 | 31.6 ± 18.7 | 0.39 | 16.3 ± 4.1 | 0.20 | 24.0 ± 9.4 | 0.35 | 0.59 |
| LA reservoir strain (%) | 16.0 ± 3.3 | 11.6 ± 2.9 | 0.13 | 10.7 ± 1.1 | 0.06 | 10.1 ± 3.5 | 0.79 | 0.59 |
| RA area ($cm^2$) | 7.6 ± 1.7 | 8.7 ± 2.4 | 0.15 | 6.6 ± 0.3 | 0.42 | 7.2 ± 0.1 | 0.14 | 0.38 |
| Interventricular septum (mm) | 9.4 ±1.3 | 8.6 ± 1.4 | 0.24 | 8.6 ± 0.3 | 0.38 | 9.2 ± 1.2 | 0.61 | 0.58 |
| Posterior wall (mm) | 7.7 ± 1.3 | 7.9 ± 1.2 | 0.72 | 7.9 ± 0.3 | 0.82 | 9.2 ± 1.0 | 0.29 | 0.22 |
| LV end-diastolic diameter (cm) | 3.9 ± 0.4 | 4.2 ± 0.7 | 0.17 | 3.5 ± 0.3 | 0.24 | 3.7 ± 0.5 | 0.50 | 0.40 |
| LV end-systolic diameter (cm) | 2.9 ± 0.2 | 3.7 ± 0.8 | 0.06 | 2.6 ± 0.3 | 0.23 | 3.3 ± 0.4 | **0.04** | 0.52 |
| LV mass (g) | 105 ± 29 | 102 ± 19 | 0.80 | 78 ± 11 | 0.23 | 89 ± 15 | 0.12 | 0.38 |
| LVEF (%) | 52 ± 8 | 23 ± 7 | **0.03** | 44 ± 5 | 0.27 | 24 ± 4 | **0.04** | 0.83 |
| LV GLS (%) | -13.4 ± 1.6 | -6.5 ± 2.0 | **0.02** | -12.8 ± 3.1 | 0.76 | -6.1 ± 1.6 | 0.07 | 0.80 |
| Average E/e’ | 6.9 ± 0.6 | 9.4 ± 2.7 | 0.18 | 9.6 ± 3.3 | 0.17 | 6.7 ± 0.6 | 0.27 | 0.19 |
| RV end-diastolic area ($cm^2$) | 9.4 ± 0.8 | 8.8 ± 1.9 | 0.60 | 7.3 ± 0.8 | **0.01** | 7.2 ± 1.1 | 0.93 | 0.29 |
| TAPSE (cm) | 1.0 ± 0.1 | 0.7 ± 0.2 | **0.04** | 1.1 ± 0.1 | 0.14 | 0.6 ± 0.1 | 0.08 | 0.53 |
| RV s’ (cm/s) | 9.2 ± 2.7 | 5.1 ± 1.4 | **0.04** | 10.1 ± 5.0 | 0.79 | 4.2 ± 0.8 | 0.19 | 0.41 |

Values are mean ± standard deviation. P values are paired t-test between baseline and terminal, or unpaired t-test between control and NOX2 shRNA.

LA — left atrial; min. — minimum; max. — maximum; RA – right atrial; LV — left ventricular; LVEF — LV ejection fraction; LV GLS — LV global longitudinal peak strain; E — early transmitral flow velocity by pulsed wave Doppler; e’ — early diastolic mitral annulus velocity by spectral tissue Doppler; RV — right ventricular; TAPSE— tricuspid annular plane systolic excursion by Mmode; RV s'— peak systolic velocity of tricuspid annulus by spectral tissue Doppler.

**Table S2**. qRT-PCR primer sequences.

| Primer name | Primer sequence |
|---|---|
| NOX forward | CAAGATGCGTGGAAACTACCTAAGAT |
| NOX2 reverse | TCCCTGCTCCCACTAACATCA |
| KCNJ2 forward | TGGATGCTGGTCATCTTCTGC |
| KCNJ2 reverse | AGCCTATGGTCGTCTGGGTCT |
| KCNJ3 forward | AGCTTCAAAAGATGGCTGGA |
| KCNJ3 reverse | TGCATATGTGACTGGGGAGA |
| KCNJ5 forward | GAAGCTCTGCCTCATGTTCC |
| KCNJ5 reverse | GTGTCGAAGCCCACGTTAAT |
| KCNJ12 forward | GGTGATCTTCAGGGGTTTGA |
| KCNJ12 reverse | CTAGAAGACCCTGCGAGGTG |
| CACNA1C forward | ACGCCTGTGACATGACCATA |
| CACNA1C reverse | GCTCTTCCTCGCTCTTCTGA |
| TBP forward | TGTATCTACAGTGAATCTTGGCTG |
| TBP reverse | GGTTCGGGGCTCTCTTATTCTC |
